# The population genomics of structural variation in a songbird genus

**DOI:** 10.1101/830356

**Authors:** Matthias H. Weissensteiner, Ignas Bunikis, Ana Catalán, Kees-Jan Francoijs, Ulrich Knief, Wieland Heim, Valentina Peona, Saurabh D. Pophaly, Fritz J. Sedlazeck, Alexander Suh, Vera M. Warmuth, Jochen B.W. Wolf

## Abstract

Structural variation (SV) accounts for a substantial part of genetic mutations segregating across eukaryotic genomes with important medical and evolutionary implications. Here, we characterized SV across evolutionary time scales in the songbird genus *Corvus* using *de novo* assembly and read mapping approaches. Combining information from short-read (*N* = 127) and long-read re-sequencing data (*N* = 31) as well as from optical maps (*N* = 16) revealed a total of 201,738 insertions, deletions and inversions. Population genetic analysis of SV in the Eurasian crow speciation model revealed an evolutionary young (~530,000 years) *cis*-acting 2.25-kb retrotransposon insertion reducing expression of the *NDP* gene with consequences for premating isolation. Our results attest to the wealth of SV segregating in natural populations and demonstrate its evolutionary significance.

---

Mutations altering the structure of DNA have the potential to drastically change phenotypes with medical and evolutionary implications (*1*–*3*). Yet, technological constraints have long impeded genome-wide characterization of (*4*). The detection of SV requires highly contiguous genome assemblies accurately representing the repetitive fraction of genomes which is known to be a vibrant source and catalyst of SV (*5*). Moreover, SV likely remains hidden unless sequence reads traverse it completely (*6*, *7*). As a consequence, despite the rapidly increasing number of short-read (SR) based genome assemblies (*8*) and associated population genomic investigations (*9*), SV generally remains unexplored. Even in genetic model organisms, population-level analysis of SV has been restricted to pedigrees (*10*) or organisms with smaller, less complex genomes (*11*, *12*), and few studies have provided a comprehensive account of SV segregating in natural populations (*12*, *13*).

To investigate the dynamics of SV and uncover its role in causing phenotypic differences, we first generated high-quality phased *de novo* genome assemblies combining long-read (LR) data from single-molecule, real-time (SMRT, PacBio) sequencing and nanochannel optical mapping (OM) for the hooded crow (*Corvus (corone) cornix*; data from (*14*)), and the European jackdaw (*Corvus monedula*). For the former, we also generated chromatin interaction mapping data (Hi-C) to obtain a chromosome-level reference genome (**Fig. 1A**, see **Supplementary Table S1** for assembly statistics). In addition, we included a previously published LR assembly of the Hawaiian crow (*Corvus hawaiiensis*) in the analyses (*15*). All assemblies were generated with the diploid-aware FALCON-UNZIP assembler (*16*), facilitating the comparison of haplotypes within species to identify heterozygous variants and determine genetic diversity at the level of single individuals. After aligning the two haplotypes of each assembly, we identified single-nucleotide polymorphisms (SNPs), insertions and deletions in all three species (**Table 1**). Genome-wide numbers of SV and SNPs per 1 Mb window were highest in jackdaw and lowest in the highly inbred Hawaiian crow (**Fig. 1B**), consistent with a positive correlation between census population size and genetic diversity (*15*, *17*).

**Table 1.**
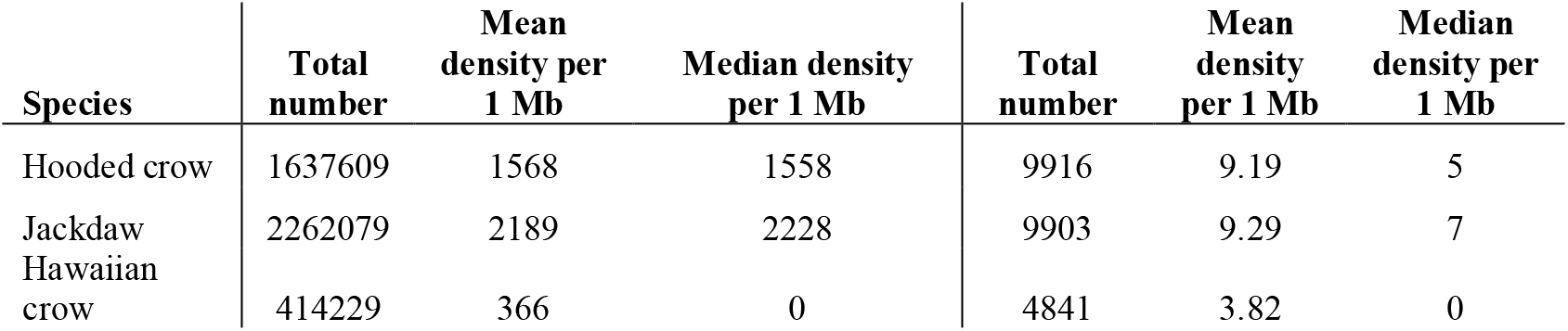
Assembly-based structural variation and single-nucleotide polymorphism detection.

| Species | Total number | Mean density per 1 Mb | Median density per 1 Mb | Total number | Mean density per 1 Mb | Median density per 1 Mb |
| --- | --- | --- | --- | --- | --- | --- |
| Hooded crow | 1637609 | 1568 | 1558 | 9916 | 9.19 | 5 |
| Jackdaw | 2262079 | 2189 | 2228 | 9903 | 9.29 | 7 |
| Hawaiian crow | 414229 | 366 | 0 | 4841 | 3.82 | 0 |

**Fig. 1.**
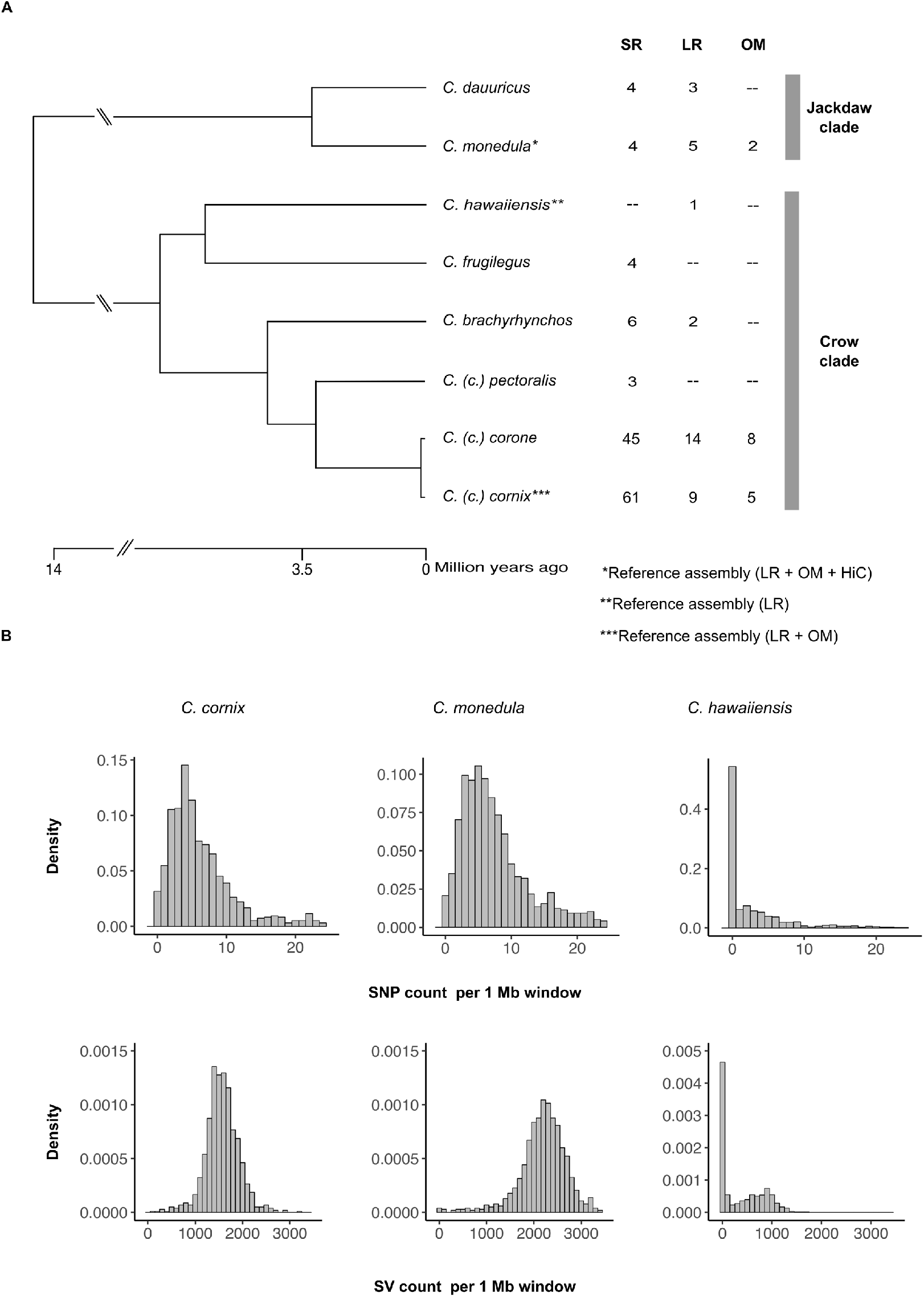
Sampling setup and assembly-based structural and single-nucleotide variation. **(A),** Phylogeny of sampled species in the genus *Corvus* (after (*50*)). Numbers in columns represent individual numbers for short-read sequencing (SR), long-read sequencing (LR) and optical mapping (OM). **(B),** Density histogram showing the abundance of genetic variation within single individuals. Counts of variants per 1 Mb windows are based on comparing the two haplotypes of each assembly. The upper panel reflects structural variation (SV) densities, the lower panel reflects densities for single-nucleotide polymorphisms (SNP).

Next, to uncover SV segregating within and between natural populations, we generated LR re-sequencing data for 31 individuals. Spanning the phylogeny of the genus, this dataset included samples from the European and Daurian jackdaw (*C. monedula, C. dauuricus*), the American crow (*C. brachyrhynchos*) and the Eurasian crow complex (*C. (corone)* spp.). The latter comprised individuals from the phenotypically divergent hooded crow (Sweden and Poland), and carrion crow populations (Spain and Germany) (*18*) (**Fig. 1A**). Individuals were sequenced to a mean sequence coverage of 15 (range: 8.47 – 27.91) with a mean read length of 7,535 bp (range: 5,219 - 10,034 bp; **Supplementary Table S2**). Mapping reads to the hooded crow reference allowed us to identify variants and genotypes for each diploid individual, which resulted in a set of 47,346 variants. SV genotyping is nontrivial and associated with high uncertainty (*7*). Thus, we utilized the sampling scheme to filter for variants complying with basic population genetic assumptions (**Fig. 2A**)(*19*). Variants that were excluded according to these criteria were enriched for deletions and clustered near the end of chromosomes (linear model, p = 10^−16^, **Fig. 2B, C**). Increased densities of repetitive elements (**Fig. 2D**), particularly tandem repeats, in these regions are conducive to erroneous genotype calling, though it is possible that a subset of these phylogenetically recurring variants indeed represent true positive, hypermutable sites.

**Fig. 2.**
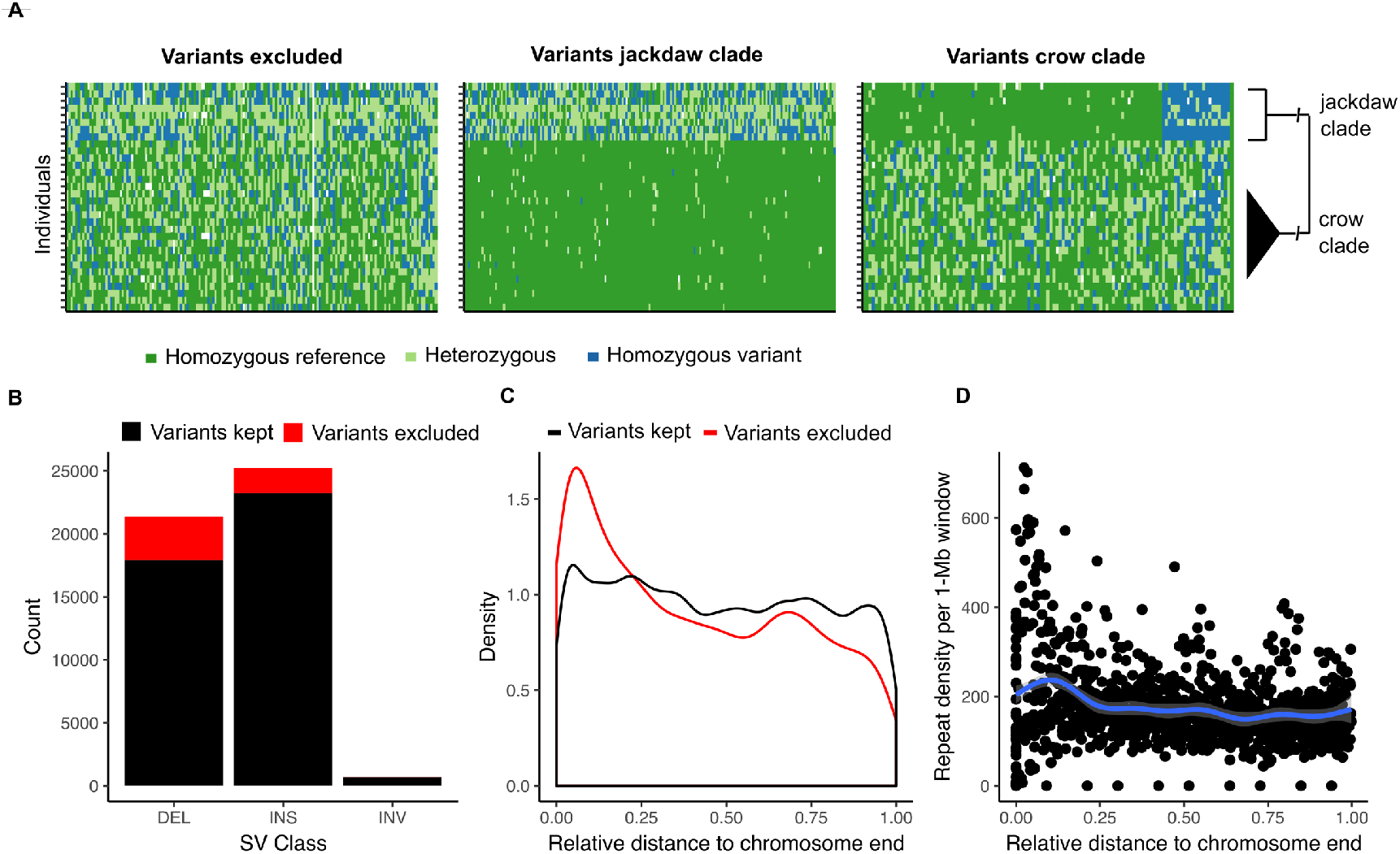
Phylogenetic filtering of read mapping-based structural variants. **(A),** Example genotype plots of LR-based variants according to phylogenetically informed filtering. Given the large divergence time of 13 million years (*50*) between the crow and jackdaw lineage, the proportion of polymorphisms shared by descent is negligible (*51*)and therefore likely constitutes false positives or hypermutable sites (left panel). Variants segregating exclusively in the jackdaw or crow clade (middle and right panel), however, comply with the infinite sites model and were retained accordingly. Plotted are genotypes of one representative chromosome (chromosome 18), with genotypes of variants in different colors, where each row corresponds to one individual (*N* = 8 individuals jackdaw clade and *N* = 24 individuals crow clade). Note that, due to the tolerance of a certain number of mis-genotyped variants per clade, some variants are present in both clades. **(B),** Excluded versus retained variants in relation to SV class and chromosomal distribution. Excluded variants are enriched for deletions (LMM, p < 10^−16^) and **c,** are most abundant at chromosome ends, coinciding with **(D),** an increased repeat density.

After the phylogenetically informed filtering step, we retained a final set of 41,868 variants (88.43 % of the initial, unfiltered set) segregating within and between species. Of these, a small proportion was classified as inversions (694, 1.657 %), whereas the vast majority was attributed to insertions (23,235, 55.495 %) and deletions (17,939, 42.846 %) relative to the hooded crow reference. Variant sizes were largest for inversions, with a median size of 980 bp (range: 51 – 99,824 bp), followed by insertions (248 bp, range: 51 – 45,373 bp) and deletions (154 bp, range: 51 – 94,167 bp). The latter showed showed noticeable peaks in the size distribution at around 900, 2,400 and 6,500 bp (**Fig. 3A**, for inversions see **Supplementary Fig. S1**), which likely stem from an overrepresentation of paralogous repeat elements. The five most common repeat motifs in insertions and deletions belonged to endogenous retrovirus-like LTR retrotransposon families and accounted for 22.78 % of all matches to a manually curated repeat library (**Supplementary Table S3**). This suggests recent activity of this transposable element group, as has been previously reported in other songbird species (*20*). More than half of all insertions and deletions could not be associated with any known repeat motif (52.19 %). The remainder was distributed approximately equally between tandem repeats (e.g. simple and low complexity repeats) and interspersed repeats. The latter category was dominated by LTR and LINE/CR retrotransposons with only a small number of SINE retrotransposons (**Fig. 3B, Table 2**). These different types of repeat elements exhibit fundamentally different mutation mechanisms (*21*) and effects on neighboring genes (*22*), such that repeat annotations are of crucial importance for the downstream population genetic analysis of SV.

**Table 2.** Characterization of LR insertions and deletions.

| Classification | Number | Percentage |
| --- | --- | --- |
| **Tandem repeat total** | 8847 | 21.48 |
| Simple repeat | 7167 | 17.4 |
| Low complexity repeat | 1595 | 3.87 |
| Satellite | 75 | 0.18 |
| rRNA | 5 | < 0.05 |
| tRNA | 3 | < 0.05 |
| Macrosatellite | 1 | < 0.05 |
| **Interspersed repeat total** | 10828 | 26.3 |
| LTR retrotransposon | 9692 | 23.53 |
| LINE / CR1 retrotransposon | 1123 | 2.27 |
| SINE retrotransposon | 11 | < 0.05 |
| D/hAT-Charlie element | 2 | < 0.05 |
| **No match** | 21492 | 52.19 |

**Fig. 3.**
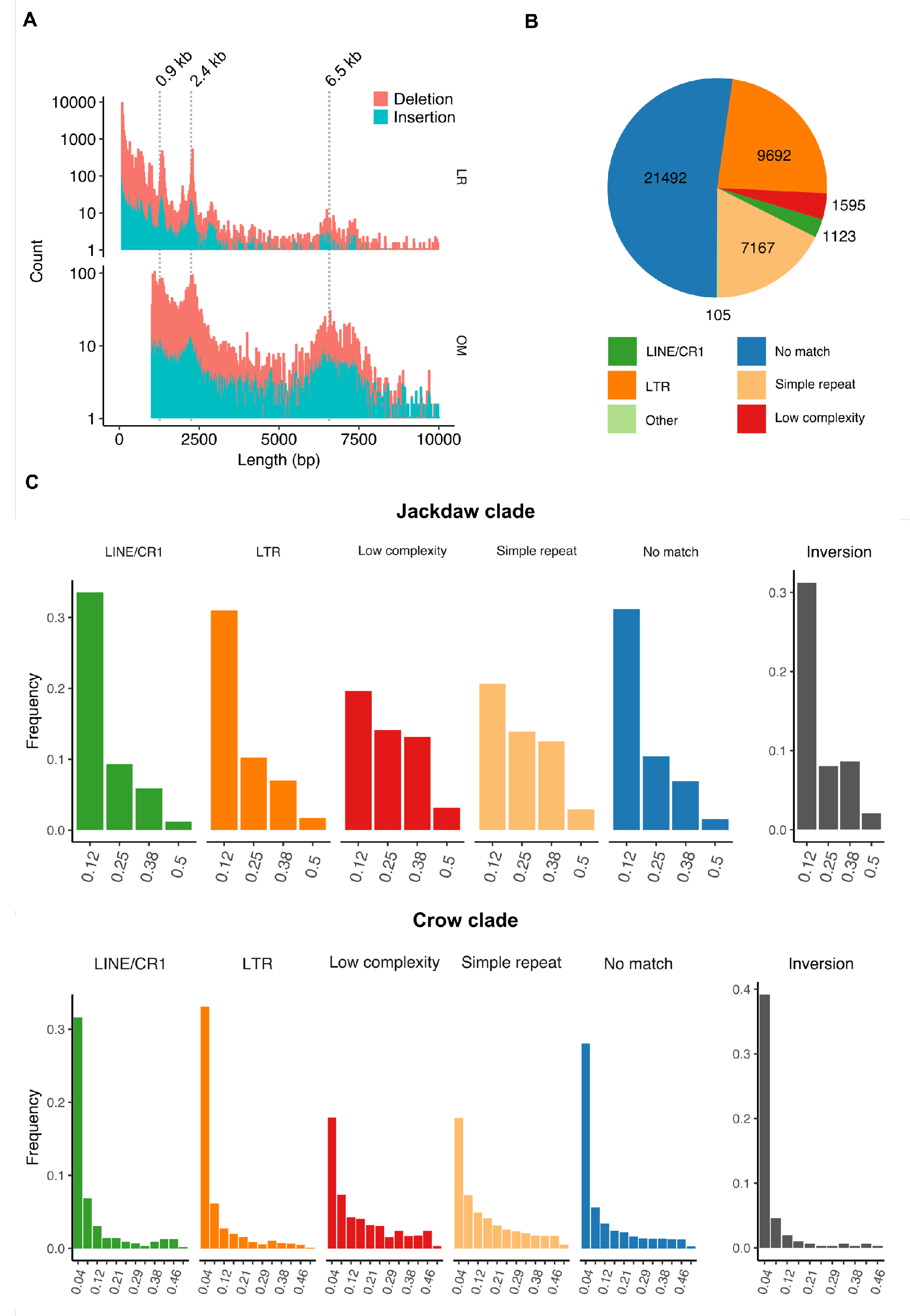
Characterization and allele frequencies of SV. **(A)** Length distributions of deletions and insertions shorter than 10 kb identified with LR (upper panel) and OM (lower panel) data. Pronounced peaks at 0.9, 2.2 kb in the LR and at 2.3 and 6.5 kb in the OM variants likely stem from an overrepresentation of specific repeats. Indeed, among the five most common repeats found in insertions and deletions are LTR retrotransposons with a consensus sequence length of 670, 1,315, 6,022 bp, respectively. **(B)** Content of insertion and deletion sequences. About half of all variants were assigned to a known repeat family, of which transposable elements from the LTR retrotransposon subclass were most common, followed by simple repeats (including microsatellites) and low complexity repeats. **(C)** Folded allele frequency spectra of structural variants. Upper and lower panels correspond to the jackdaw and crow clade, respectively. The five left panels depict the minor allele frequencies of insertions and deletions, and the rightmost panel that of inversions.

We then scrutinized structural variation segregating within clades sharing recent common ancestry. A total of 35,723 and 29,555 variants remained after filtering in the jackdaw (*N* = 8 individuals; *C. monedula, C. dauuricus*) and crow clade (*N* = 24; *C. (corone) spp., C. brachyrhynchos*), respectively. Using the full data set across all populations within each clade allowed us to compare folded allele frequency spectra between SV classes and repeat types with high resolution (for population specific spectra unbiased by population structure see **Supplementary Fig. S2**). Consistent with recent studies in grapevine and *Drosophila* SV (*12*, *23*), the distribution of allele frequencies was skewed towards rare alleles (**Fig. 3C**). However, allele frequency spectra of different SV classes differed in shape. While insertions and deletions associated with LTR elements, LINE/CR1 elements or without any known match as well as inversions exhibited the typical pattern of a strongly right-skewed frequency distribution, allele frequencies of simple and low complexity repeats were shifted towards intermediate frequencies. Besides a potential technical bias due to the more difficult genotyping and variant discovery of these classes (*24*), this pattern is consistent with convergence to intermediate allele frequencies due to high mutation rates (*21*). These results illustrate how different underlying mutation dynamics potentially impact the analysis of population genetic parameters for SV.

To improve our ability to detect larger SV and to provide an independent orthogonal approach for SV discovery, we generated an additional 14 optical maps (**Fig. 1A**) and compared them to the hooded crow reference assembly. Following that approach, we identified 12,807 insertions, 8,799 deletions and 293 inversions. As expected from the increased size of individually assessed DNA molecules (mean molecule N50 = 223.38 kb), variants identified with this approach exhibited a different size range (**Fig. 3A**) after applying the same upper limit (100 kb) as for the LR SV calls and a lower limit of resolution (1 kb) (*25*). Interestingly, insertion and deletions were not only enriched at lengths around 0.9 and 2.4 kb as seen in the LR-based SV calling, but also at ~ 6.5 kb, indicating an influence of the TguERV1-Ld_I_corCor LTR retrotransposon, which was the third most common single repeat in the LR variant set with a consensus sequence length of 6,022 bp (**Supplementary Table S3**). Thus, independent approaches targeting different size ranges of SV are vital to increase sensitivity in detecting hidden genetic variation.

To increase our sample size and expand our analysis to further populations and species (**Fig. 1A**), we applied a combination of three different short-read (SR) based SV detection approaches on previously published data of 127 individuals (*18*, *26*). In total, we identified 132,025 variants of which 97,524 (73.87 %) were unique to single individuals. In total, only 11,951 variants overlapped with the final set of variants identified in the long-read data set (corresponding to 9.05 % of SR and 28.54 % LR calls). This disconnect cannot be explained solely by differences in sample size. More likely, it indicates a high number of false-positives and false-negatives in the SR-based approach known for its sensitivity to the calling method (*27*) and disparity to LR-based calls (*7*). Therefore, we focused on the LR-based SV calls in the subsequent analysis and considered SR calls only for specific mutations.

Next, we investigated population structure using principal component analyses (PCA). The pattern in **Fig. 4A** (based on LR data) recapitulates the pattern of population stratification found in Vijay et al. based on 16.6 million SNPs (*18*), and thus supports the general suitability of SV genotypes for population genetic analyses (for SR data see **Supplementary Fig. S3**). In order to identify SV associated with prezygotic reproductive isolation, we calculated genetic differentiation between phenotypically divergent populations connected by gene flow (*18*, *26*) and allopatric populations within the same phenotype (*18*). Mean F_ST_ was low overall with values ranging from 0.03 in the hooded versus carrion crow comparison to 0.156 in the hooded versus American crow comparison.

**Fig. 4.**
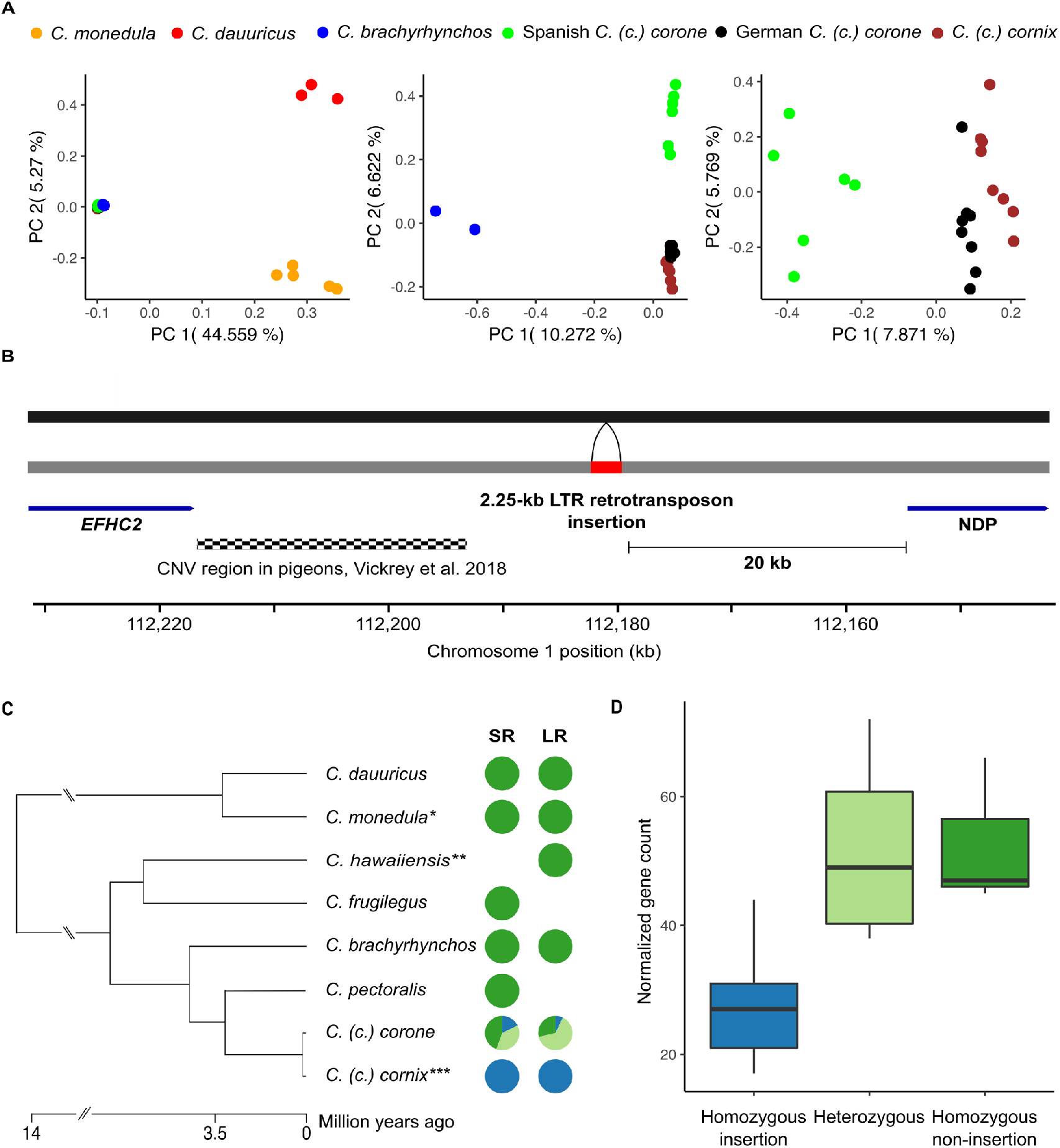
SV-based population structure and LTR retrotransposon insertion upstream of the *NDP* gene. **(A)** Principal component analysis based on SV genotypes. The first two principal components separate the crow and jackdaw clade, while principal components 3 to 5 separate lineages within the crow clade. **(B)** A 2.25-kb LTR retrotransposon insertion into the crow lineage (black bar: ancestral state, grey bar: derived, reference allele) belongs to the endogenous retrovirus-like family ERVK and the subfamily TguERV1-Ld-I and is located 20 kb upstream of the *NDP* gene. In close proximity, variation in copy number is associated with plumage pattern variation in pigeons. **(C)** Genotypes of the LTR element in short-read (SR) and long-read (LR) data. In both datasets, the LTR element insertion (blue) is fixed in all hooded crow populations. Species and populations with a black plumage are either polymorphic (light green) or fixed non-insertion (green). **(D)** Gene expression of *NDP*. Normalized gene counts of 18 individuals are significantly associated with the insertion genotypes (LMM, p = 0.002).

A total of 103 variants fell into the 99^th^ percentile of F_ST_ in the gray-coated hooded versus all-black carrion crow population comparison in central Europe. These variants, located on in total 23 chromosomes, were considered as *ad hoc* candidate outlier loci subject to divergent selection (*9*), and were found at a median distance of 14.32 kb to adjacent genes (range: 0 - 695.84 kb). (**Supplementary Table S4**). Ten of these outliers (10.31 %) were placed on chromosome 18, which only represents 1.22 % of the entire assembly, corresponding to an ~8.5-fold enrichment. Given that outliers are located in the proximity of previously identified genes presumably under divergent selection (such as *AXIN2* and RGS9, Supplementary Table S5), this supports a crucial role of chromosome 18 in maintaining plumage divergence (*26*, *28*).

The three highest F_ST_ outliers included an 86 bp indel on chromosome 18 inside of a tandem repeat array, a 1.56 kb indel on chromosome 3 and a 2.25 kb indel on chromosome 1 (**Supplementary Table S5**). The latter, an LTR retrotransposon insertion, was located 20 kb upstream of the *NDP* gene on chromosome 1 (**Fig. 4B)**, a gene known to contribute to the maintenance of color divergence across the European crow hybrid zone (*28*). Molecular dating based on the LTR region suggest an insertion event at <534,000 years ago upon diversification of the European crow lineage (**Fig. 4C**) (*18*). In current day populations, the insertion still segregates in all-black crows including *C (c.) corone* in Europe and *C (c.) orientalis* in Russia (*N* individuals with LR = 14 and with SR = 45 genotypes) (**Fig. 4C)**. All hooded crow *C. (c.) cornix* individuals, however, genotyped with LR (*N* = 9) and SR data (*N* = 61) were homozygous for the insertion regardless of their population of origin. This finding is consistent with a selective sweep in proximity to the *NDP* gene that has previously been suggested for hooded crow populations (*26*, *28*). Recent work has also shown that the *NDP* gene exhibits decreased gene expression in grey feather follicles of hooded crows, suggesting a role in modulating overall plumage color patterning (*29*). Following re-analysis of normalized gene expression data for 8 carrion and 10 hooded crows (*29*), we found a significant association between the homozygous insertion genotype and decreased *NDP* gene expression levels (linear model, p = 0.002) (**Fig. 4D**), consistent with reduced pigmentation in hooded crows (*29*).

To further investigate the relationship between the abovementioned insertion and phenotypic differences between all-black *C (c.) corone* and gray-coated *C. (c.) cornix* populations, we genotyped 120 individuals from the European hybrid zone using PCR (Methods, (*28*)). Including data of adjacent SNPs for the same individuals, we tested the association between genotype and pigmentation phenotype. A statistical model including the insertion fit best to the observed phenotypes (ΔAICc = 2.33, but ΔBIC = −0.12) explaining an additional 10.32% of the variance of the phenotype-derived PC1 relative to the adjacent SNPs. The insertion lies upstream of *NDP* in close proximity to an orthologous region in pigeons containing a copy number variation shown to modulate plumage patterning (**Fig. 4B**) (*30*). Reminiscent of the wing color altering TE insertion in the peppered moth (*3*), this insertion thus constitutes a prime candidate causal mutation modulating gene expression with phenotypic consequences; reminiscent of the TE insertion in the peppered moth altering wing coloration (*3*). While such insertions have usually been associated with increased expression of the affected gene (*31*), there are also examples of TE insertions repressing gene activity, as observed here (*32*).

In conclusion, this study provides the first comprehensive population-level SV catalogue in a non-model organism, further elucidating the role of SV on modulating expression of evolutionary important genes with phenotypic consequences. Given that the majority of SV is likely still uncovered in most organisms (*33*), these results mark an important hallmark for the field highlighting the evolutionary importance of SV in natural populations and the need for rigorous methodological approaches.

## Material and Methods

### Short-read sequencing data

We compiled raw short-read sequencing data from Poelstra et al. 2014 and Vijay et al. 2016 (*18*, *26*) for *Corvus (corone)* spp., *C. frugilegus, C. dauuricus*, *C. monedula* and *C. brachyrhynchos* (for more information on the origin of samples and accession numbers of the data see **Supplementary Table S5**). Overall, 127 individuals had an average 12.6-fold sequencing coverage using paired-end libraries (primarily) sequenced on an Illumina HiSeq2000 machine.

### DNA extraction and long-read sequencing

First, we extracted high-molecular weight DNA from a total of 32 samples using either a modified phenol-chloroform extraction protocol (*14*), or the Qiagen Genomic-tip kit (following manufacturer’s instructions) from frozen blood samples. For sampling details, see **Supplementary Table S5**. Extracted DNA was eluted in 10 mM Tris buffer and stored at −80 °C. The quality and concentration of the DNA was assessed using a 0.5 % agarose gel (run for >8 h at 25 V) and a Nanodrop spectrophotometer (ThermoFisherScientific). Long-read sequencing DNA libraries were prepared using the SMRTbell Template Prep Kit 1.0 (Pacific Biosciences). For each library, 10 μg genomic DNA was sheared into 20-kb fragments with the Hydroshear (ThermoFisherScientific) instrument. SMRTbell libraries for circular consensus sequencing were generated after an Exo VII treatment, DNA damage repair and end-repair before ligation of hairpin adaptors. Following an exonuclease treatment and PB AMPure bead wash, libraries were size-selected using the BluePippin system with a minimum cutoff value of 8,500 bp. All libraries were then sequenced on either the RSII or Sequel instrument from Pacific Biosciences, totaling 324 RSII and 76 Sequel SMRT cells, respectively, resulting in 754 Gbp of raw data.

### Genome assembly

In birds, females are the heterogametic sex (ZW). For this study, we were interested in a high-quality assembly of all autosomes and the shared sex chromosome (Z) and accordingly chose male individuals for the genome assemblies. Note, however, that this choice excludes the female-specific W chromosome *a priori*. Diploid genome assembly was performed for both a hooded crow and a jackdaw individual. For the former a long-read based genome assembly has previously been published (*14*) and is available under the accession number GCA_002023255.2 at the repository of the National Center for Biotechnology Information (NCBI, www.ncbi.nlm.nih.gov). Here, we (re)assembled raw reads using updated filtering and assembly software. First, all SMRT cells for the respective individuals (102 for the hooded crow individual S_Up_H32, 70 for the jackdaw individual S_Up_J01) were imported into the SMRT Analysis software suite (v2.3.0). Subreads shorter than 500 bp or with a quality (QV) <80 were filtered out. The resulting data sets were used for *de novo* assembly with FALCON UNZIP v0.4.0 (*16*). Initial FALCON UNZIP assemblies of hooded crow and jackdaw consisted of primary and associated contigs with a total length of 1,053.37 Mb and 965.95 Mb for the hooded crow and 1,073.84 and 1,092.55 Mb for the jackdaw, presumably corresponding to the two chromosomal haplotypes (for assembly statistics see **Supplementary Table S1**). To further improve the assembly, we performed consensus calling of individual bases using ARROW (*16*). In addition, we obtained the genome of the Hawaiian crow (*Corvus hawaiiensis*) from the repository of NCBI with accession number GCA_003402825.1. This genome had been likewise derived from long-reads generated with the SMRT technology and assembled using FALCON UNZIP (*16*). To assess the completeness of the newly assembled genomes we used BUSCO v2.0.1 (*34*). The aves and the vertebrate databases were used to indentify ultra conserved orthologous gene sets (**Supplementary Table S1**).

### Optical mapping data and assembly

We generated additional optical map assemblies for two jackdaw individuals, 8 carrion crow individuals and 4 additional hooded crow individuals, following the same approach used for the optical map assembly of the hooded crow individual (see Weissensteiner et al.(*14*)). In brief, we extracted nuclei of red blood cells and captured them in low-melting point agarose plugs. DNA extraction was followed by melting and digesting of the agarose resulting in a high-molecular weight DNA solution. After digestion with a nicking endonuclease (Nt.BspQI) which inserts a fluorescently labelled nick strand, the sample was loaded onto an IrysChip, which was followed by fluorescent label detection on the Irys instrument. The assembled consensus maps were then used to perform SV calling as part of the Bionano Access 1.3.1 Bionano Solve pipeline 3.3.1 (pipeline version 7841). As reference an *in-silico* map of the hooded crow reference assembly was used. Molecule and assembly statistics of optical maps can be found in **Supplementary Table S6**. For details regarding the hybrid scaffolding see Weissensteiner et al. 2017 (*14*).

### Hi-C chromatin interaction mapping and scaffolding

One Dovetail Hi-C library was prepared from a hooded crow sample following Lieberman-Aiden et al. (2009) (*35*). In brief, chromatin was fixed in place with formaldehyde in the nucleus and extracted thereafter. Fixed chromatin was digested with DpnII, the 5’ overhangs filled with biotinylated nucleotides and free blunt ends were ligated. After ligation, crosslinks were reversed and the DNA purified from the protein. Purified DNA was treated such that all biotin was removed that was not internal to ligated fragments. The DNA was then sheared to ~350 bp mean fragment size and sequencing libraries were generated using NEBNext Ultra enzymes and Illumina-compatible adapters. Biotin-containing fragments were isolated using streptavidin beads before PCR enrichment of each library. The library was then sequenced on an Illumina HiSeq X (rapid run mode). The Dovetail Hi-C library reads and the contigs of the primary FALCON UNZIP assembly were used as input data for HiRise, a software pipeline designed specifically for using proximity ligation data to scaffold genome assemblies (*36*). An iterative analysis was conducted. First, Hi-C library sequences were aligned to the draft input assembly using a modified SNAP read mapper (http://snap.cs.berkeley.edu). The separation of read pairs mapped within draft scaffolds were analyzed by HiRise to produce a likelihood model for genomic distance between read pairs, and the model was used to identify and break putative misjoins, to score prospective joins and make joins above a threshold. The resulting 48 super-scaffolds were assigned to 27 chromosomes based on synteny to the flycatcher genome version (NCBI accession GCA_000247815.2) (*37*) using LASTZ (*38*). The final Hi-C scaffolded hooded crow assembly is available as a Dryad repository, file XX.

### Assembly-based SV and SNP detection

We aligned the associated contigs of all three assemblies (hooded crow, jackdaw and Hawaiian crow) to the primary contigs (super-scaffolded to chromosome level for hooded crow) using MUMmer (*39*). SNPs were then identified using *show-snps* with the options –Clr and –T following a filtering step with delta-filter –r and –q. We only considered single-nucleotide differences in this analysis.

Structural variants between the two haplotypes of each assembly were identified using two independent approaches. First, we used the alignments produced with MUMmer to identify variants using the Assemblytics tool (*40*). We then converted the output to a *vcf* file using SURVIVOR (v1.0.3) (*27*). Independently, we used the smartie-sv pipeline to identify structural variants (*41*), and then converted and merged the output with the Assemblytics-based variant set with SURVIVOR. This final unified variant set was then used to calculated SV-density in non-overlapping 1-Mb windows.

### Repeat annotation and characterization of insertions and deletions

To characterize the repeat content of the hooded crow assembly, we used the repeat library from Vijay et al. (*18*). Raw consensus sequences were manually curated following the method used in Suh et al. (2018) (*20*). Every consensus sequence was aligned back to the reference genome, then the best 20 BLASTN (*42*) hits were collected, extended by 2 kb and aligned to one another using MAFFT (v6; (*43*)). The alignments were manually curated applying a majority-rule and the superfamily of each repeat assessed following Wicker et al. (*44*). We then masked the new consensus sequences in CENSOR (http://www.girinst.org/censor/index.php) and named them according to homology to known repeats present in Repbase(*45*). Repeats with high sequence similarity to known repeats were given the name of the known repeat + suffix “_corCor”; repeats with partial homology were named with the suffix “-L_corCor” where “L” stands for “like” (*20*). Repeats with no homology to other known repeats were considered as new families and named with the prefix “corCor” followed by the name of their superfamilies. Using this fully curated repeat library (**Supplementary file S1**), we performed a RepeatMasker (*46*) search on all sequences reported for insertion and deletion variants. In case of multiple different matches per variant or individual, we took the match with the highest overlap with the query sequence to yield a single match for each variant. We also performed a RepeatMasker search with the curated library to estimate repeat density per 1 Mb window in the hooded crow reference assembly.

### Read-mapping based SV and SNP detection

We aligned PacBio long-read data of all re-sequenced individuals to both the hooded crow and jackdaw reference assembly using NGM-LR (*47*) (v0.2.2) with the –pacbio option and sorted and indexed resulting alignments with samtools (*48*)(v1.9). Initial SV calling per individual was then performed using Sniffles (*47*) (v1.0.8) with parameters set to a minimum support of 5 reads per variant (--min_support 5) and enabled – genotype, –cluster and –report_seq options. We removed abundant translocation calls indicative of an excess of false positives and filtered remaining variants for a maximum length of 100 kb and a maximum read support of 60 with bcftools (*49*). Both of these filtering steps have been shown to be necessary to remove erroneously called variants. Next, we generated a merged multi-sample *vcf* file consisting of all individuals from both the crow and the jackdaw clades with SURVIVOR merge and options set to 1000 1 1 0 0 50. This merged *vcf* file was then used as an input to reiterate SV calling with Sniffles for each individual with the –Ivcf option enabled, effectively genotyping each variant per individual. Resulting single individual *vcf* files were again merged with the SURVIVOR command described above and variants overlapping with assembly gaps were removed. We converted the *vcf* file into a genotype file with vcftools (*49*) (v0.1.15) for downstream analysis.

To account for the high amount of genotyping errors and false positives after initial filtering, we employed a ‘phylogenetic’ filtering strategy. The crow and jackdaw clades diverged roughly 13 million years ago (*50*), such that the proportion of polymorphisms shared by descent is near negligible (*51*). Moreover, under the infinite sites model, recurrent mutations are not expected, such that polymorphisms segregating in both lineages most likely constitute false positives. For population genetic analyses of the jackdaw clade, we therefore considered only variants which were homozygous for the reference in crow clade individuals, allowing for a maximum of four genotyping errors. In the crow clade analyses, we only retained variants which were either fixed for the reference or the variant allele in the jackdaw clade, allowing for 2 genotyping errors. It is likely that this conservative approach excludes variants with a high mutation rate (*52*). However, since it is difficult to differentiate such variants from genotyping errors, we deemed this filter necessary to yield a set of more reliable variants. Due to the tolerance of genotyping errors, there is a number of variants present in both clades, most of them fixed or almost fixed in both clades. Extensive manual curation would be necessary to differentiate between genotyping errors and variants truly polymorphic between clades. To find common features in filtered versus kept variants, we applied a generalized linear mixed-effects model with a binomial error structure, in which we coded the dependent variable as 1 for a retained variant and as 0 for a filtered variant. As covariates we included the distance to the chromosome end and variant class as a factor (insertion, deletion or inversion). We further fitted chromosome identity as a random intercept term. All models were run in R (v3.2.3, R Core Team) using the lme4 package (*53*) (v1.1-19).

The short-read data were mapped using BWA-MEM with the –M option to the hooded crow reference assembly (*54*). We used LUMPY (*55*), DELLY (*56*) and Manta (*57*) to obtain SV calls for each sample using their respective default parameters. Subsequently the individual SV calls per sample were merged using SURVIVOR (*27*) merge with the parameters: “1000 2 1 0 0 0”. This filtering step retained only SV calls for which 2 out of the 3 callers had reported a call within 1 kbp. Next, we computed the coverage of low mapping quality reads (MQ<5) for each sample independently and recorded regions where the low MQ coverage exceeded 10. SV calls which overlapped these regions were filtered out.

### Optical mapping-based SV detection

The assembled optical maps were used to identify SV compared to the provided reference, which is part of the assembly pipeline or can be run manually. SV calling iwas based on the alignment between an individual assembled consensus cmap and the *in-silico* generated map of the reference using a multiple local alignment algorithm and detecting SV signatures. The detection algorithm identifies insertions, deletions, translocation breakpoints, inversion breakpoints and duplications. The results are in a generated file in the Bionano specific format smap in which the SVs are classified as homozygous or heterozygous. This resulting smap file was converted to vcf format (version 4.2) for further downstream processing.

### Population genetic analysis of structural variants

To investigate population structure, we performed principal component analyses (PCA) with both the long-read and short-read variant sets using the R packages SNPrelate (v1.4.2.) and gdsfmt (v1.6.2) (*58*). We further calculated the folded allele frequency spectrum using minor allele frequencies of variants for all populations and clades. To estimate genetic differentiation of structural variations, we calculated F_ST_ for each variant using vcftools (*59*). We employed the Weir and Cockerham estimator for F_ST_ (*60*), variants with an F_ST_ exceeding the 99^th^ percentile were considered as outliers.

### Analyses of SV in the vicinity of the *NDP* gene

The LTR retrotransposon insertion identified upstream of the *NDP* gene on chromosome 1 - an ERV1 element belonging to the subfamily TguERV1-Ld-I - has initially been called as a deletion relative to the reference (hooded crow) assembly. To estimate its age, we assumed that the two long terminal repeats of the full-length LTR retrotransposon were identical at the time of insertion (*61*). Thus, we quantified the number of substitutions and 1-bp indels between the left and right LTR of the insertion at position 112,179,329 on chromosome 1 of the hooded crow reference. The LTRs showed 5 differences which we then divided by the length of the LTR (296 bp) and by twice the neutral substitution rate per site and million years (0.0158 (*18*)). Assuming that all differences between the left and right LTR of this insertion are fixed, this estimate yields an upper bound of the insertion age. However, overlap with SNPs segregating in the hooded crow population suggests that all 5 differences were not fixed and the insertion could thus be considerably younger.

To investigate a potential link between the LTR insertion and differences in plumage coloration, we re-analyzed gene expression data from 10 black-and-grey hooded crows and 8 all-black carrion crows raised under common garden conditions (*29*). Expression was measured for messenger RNA derived from feather buds at the torso, where carrion crows have black feathers and hooded crows are grey. We inferred the insertion genotype for each individual using short-read sequencing data via visual inspection of the alignments to the hooded crow reference. We then fitted a linear model with normalized *NDP* expression data as the dependent variable and *NDP* indel genotype as the predictor. We decomposed the effect of the insertion genotype into an additive component (the number of non-inserted minor allele copies – 0, 1, or 2 – as a covariate) and a dominance component (homozygous = 0, heterozygous = 1).

To further establish a link between the LTR retrotransposon insertion and phenotypic differences, we made use of a hybrid admixture data set from the European hybrid zone (*28*). We designed three sets of PCR primers to genotype the insertion for 120 phenotyped individuals from the European hybrid zone of all-black *C. (c.) corone* and black-and-grey *C. (c.) cornix* crows. For absence of the insertion, a pair of primers located in the sequence flanking the insertion was used (A_F_3 ‘AGTAACTGTCCTCTGTAGTGCAGG’ and A_R_3 ‘CCTGGGTAAGATCACAGTGTTGC’) resulting in a 197 bp fragment. For presence of the insertion, a pair of primers with one in the flanking and one in either left or right LTR region of the insertion (P_L_F_1 ‘TCCTCTGTAGTGCAGGACTGG’ and P_L_R_2 ‘CACCCATGGTTTCCCTCACA’, as well as P_R_F_1 ‘GGATCGGGGATCGTTCTGCT’ and P_R_R_1 ‘CACAGCCCCAGAAGATGTGC’), resulting in fragments of 659 and 564 bp, respectively. A representative gel picture used for genotyping can be found in the **Supplementary Fig. S4.** Phenotypic data was taken from Knief et al. (*28*) who summarized 11 plumage color measures on the dorsal and ventral body into a principal component (PC1), explaining 78% of the phenotypic variation. We then tested whether the interaction between chromosome 18 and the insertion genotype explained more variation in plumage color than the interaction between chromosome 18 and the most significant SNP near the *NDP* gene (*28*). We fitted two linear regression models on the same subset of the data that contained no missing genotypes (*N* = 120 individuals). In both models, we used color PC1 as our dependent variable. In the first model, we fitted the interaction between chromosome 18 and the insertion genotype, and in the second model the interaction between chromosome 18 and the SNP genotype as our independent variables. Both variables were coded as 0, 1, 2 copies of the derived allele and fitted as factors. We selected the model with the better fit to the data by estimating the AICc and BIC and deemed a ΔAICc ≥ 2 as significant.

## Acknowledgements

We thank John Marzluff, Vittorio Bagglione and Kristaps Sokolovskis for providing sample material. Reto Burri and Sergio Tusso Gomez provided helpful input for the downstream analysis. We are thankful for being able to use the UPPMAX Next-Generation Sequencing Cluster and Storage (UPPNEX) project, funded by the Knut and Alice Wallenberg Foundation and the Swedish National Infrastructure for Computing. This work was supported by the Swedish Research Council (grant number 621-2010-5553 to J.B.W.W. and grant number 2016-05139 to A.S.), the European Research Council (grant number ERCStG-336536 to J.B.W.W.) and the National Institutes of Health (grant number UM1 HG008898 to F.J.S).

## Author contributions

M.W. and J.W. conceived of the study, conducted field work and wrote the manuscript with input from all other authors. M.W. conducted lab work and all bioinformatic analyses with help from V.P., V.W., S.D.P., A.S. (repeat annotation) and U.K. (statistical analyses). W.H. conducted field work. I.B. generated genome assemblies and F.J.S. performed short-read based SV calling.

