## Supplementary Material for "The population genomics of structural variation in a songbird genus"

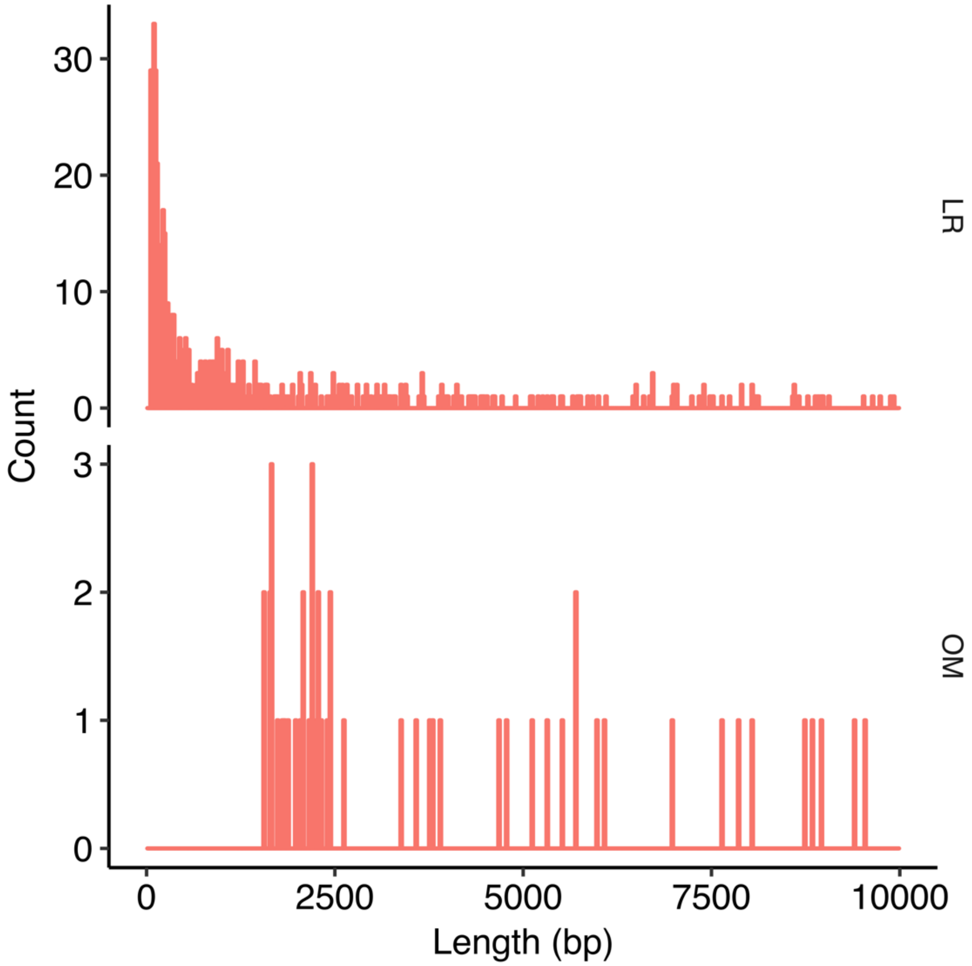


**Fig. S1.**
Length distribution of inversions shorter than 10 kb identified with LR (top panel) and OM (bottom panel).

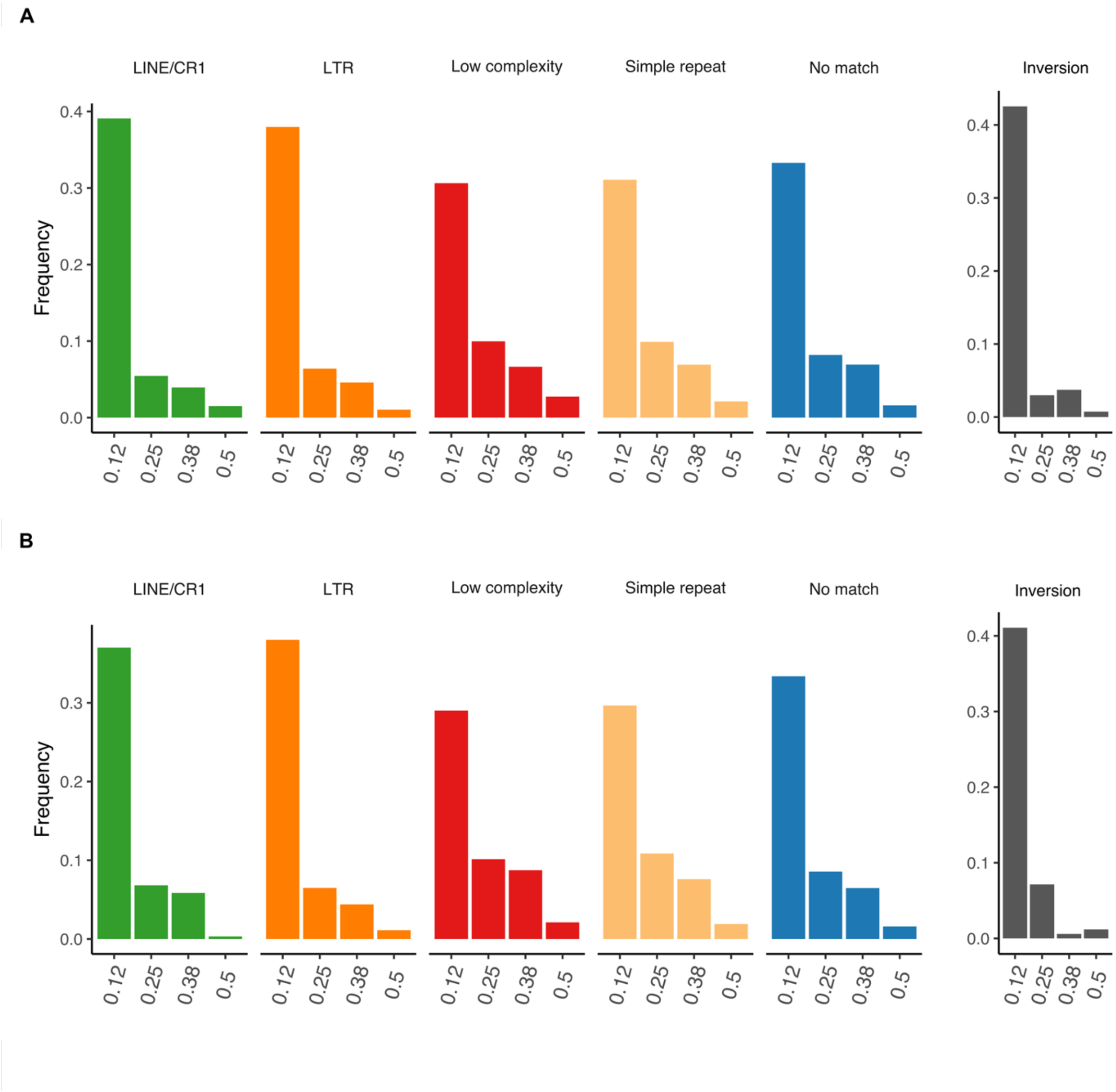


**Fig. S2.**
Folded allele frequency spectra of A, the hooded crow population and B, the German carrion crow population.

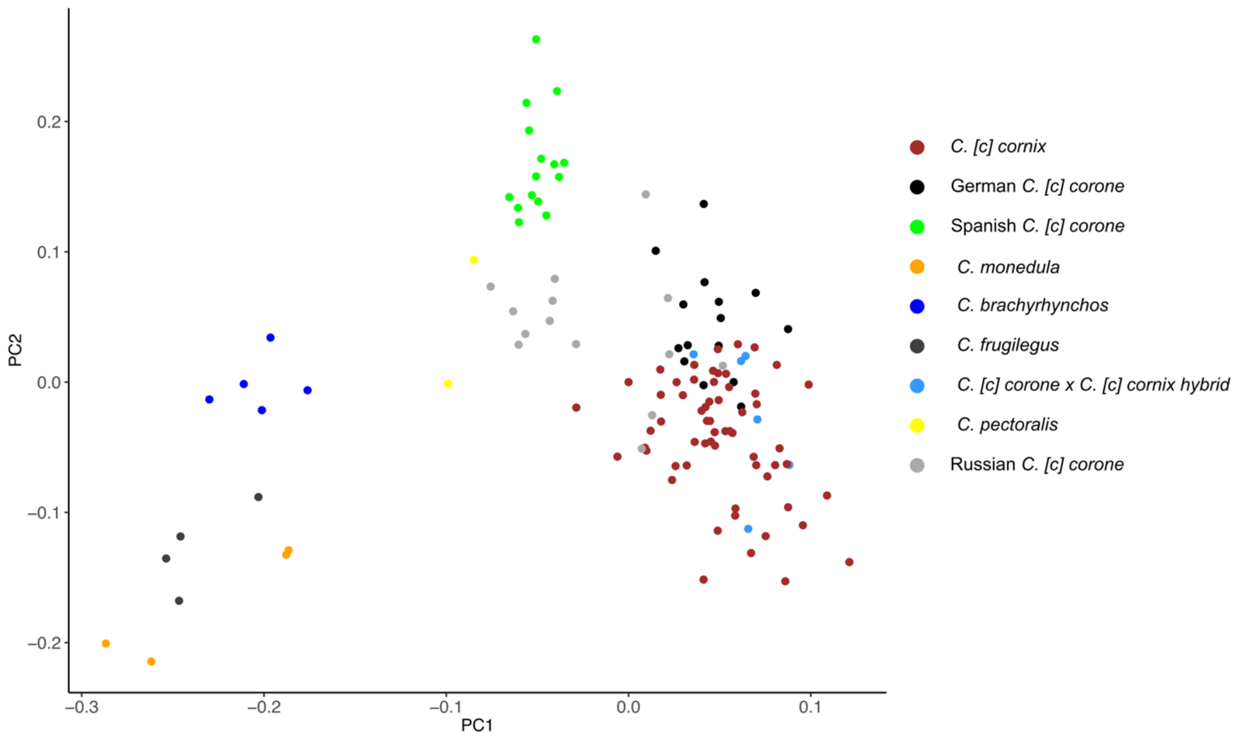


**Fig. S3.**
Principal component analysis of SV based on SR data.

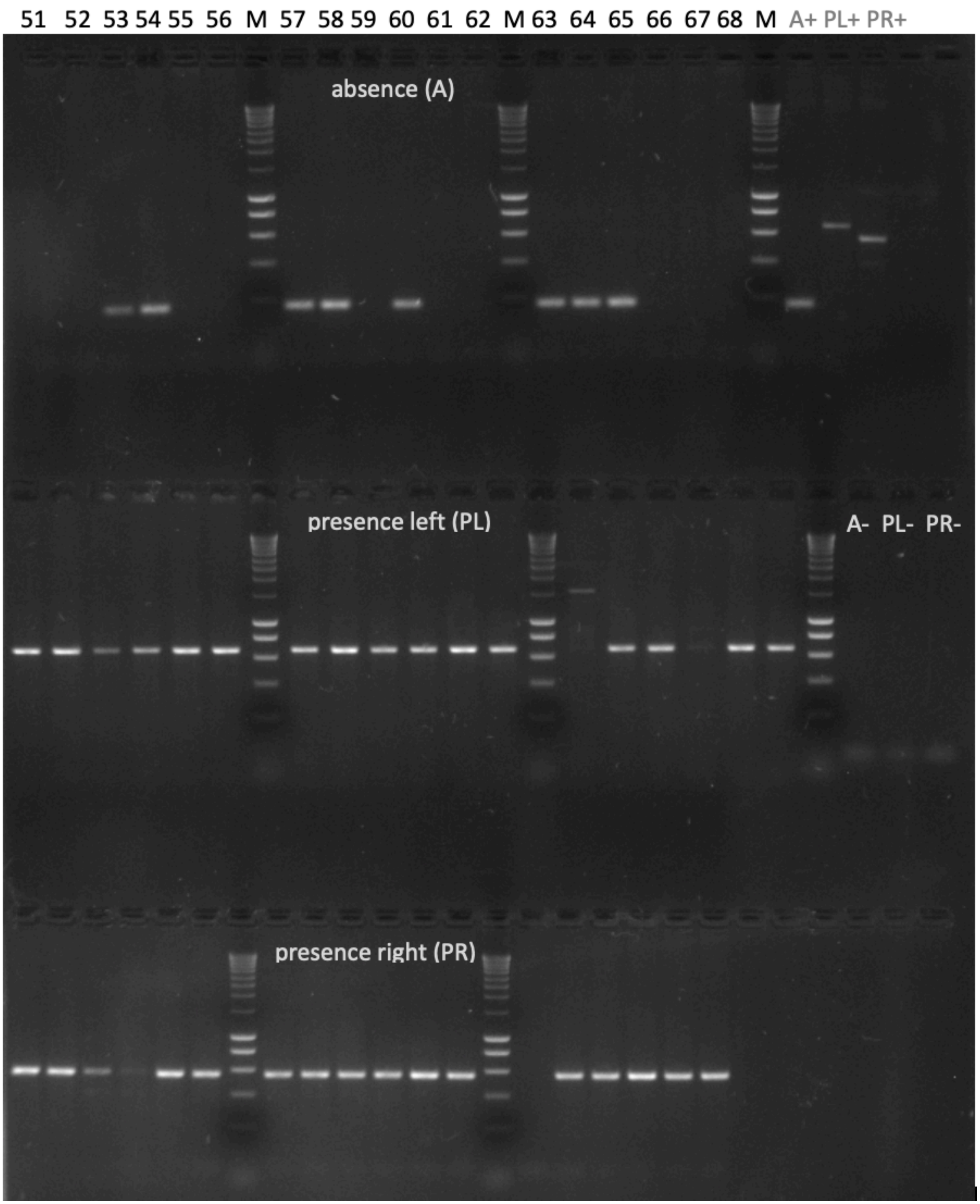


**Fig. S4.**

Representative gel picture of the LTR retrotransposon insertion genotyping in the vicinity of the *NDP* gene on chromosome 1. Numbered columns show focal individual genotyped for three different PCR fragments.

**Table S1.**

Assembly statistics of generated assemblies.

| Assembly | Number of scaffolds | Total length | Longest scaffold | Mean scaffold length | Median scaffold length | Scaffold N50 | Number of contigs | Total length | Longest contig | Contig N50 | % complete BUSCOs vertebrate | % complete BUSCOs aves |
|---|---|---|---|---|---|---|---|---|---|---|---|---|
| Hooded crow - Super-scaffolded primary assembly | 48 | 1037.32 | 154.82 | 21.61 | 6.88 | 74.11 | 810 | 1033.98 | 33.50 | 7.83 | 94,39 | 88,67 |
| Hooded crow - FALCON UNZIP associated assembly | - | - | - | - | - | - | 7645 | 965.95 | 1.80 | 0.22 | 83,49 | 78,90 |
| Jackdaw - Super-scaffolded primary assembly | 136 | 1042.43 | 58.30 | 7.66 | 3.436 | 16.38 | 1607 | 1036.86 | 52.68 | 12.14 | 93,70 | 93,70 |
| Jackdaw - FALCON UNZIP associated assembly | - | - | - | - | - | - | 6349 | 1009.69 | 3.10 | 0.42 | 84,30 | 84,30 |
| Hawaiian crow - FALCON UNZIP primary assembly | - | - | - | - | - | - | 670 | 1064.97 | 31.53 | 7.73 | 95,05 | 95,05 |
| Hawaiian crow - FALCON UNZIP associated assembly | - | - | - | - | - | - | 2082 | 432.63 | 9.52 | 0.45 | 42,65 | 39,47 |

**Table S2.**

Long-read sequencing data. Summary of sequencing data generated with PacBio SMRT-sequencing technology.

| Run_ID | Number of reads | Amount sequenced [Gbp] | Longest read [kbp] | Mean read length [kbp] | Median read length [kbp] |
|---|---|---|---|---|---|
| D_Ko_C29__pb_247_001 | 1319078 | 10,80 | 78,574 | 8,187 | 7,726 |
| E_Vi_C58__pb_375_006 | 1426456 | 13,54 | 53,867 | 9,493 | 8,682 |
| S_To_J14__pb_382_001 | 1646343 | 13,18 | 61,411 | 8,005 | 7,252 |
| S_Up_H24__pb_382_002 | 1543348 | 13,59 | 67,679 | 8,805 | 7,837 |
| S_Up_H29__pb_382_003 | 1378152 | 13,38 | 70,966 | 9,709 | 8,756 |
| USA_Wa_B02__pb_383_005 | 2282703 | 18,93 | 53,388 | 8,294 | 7,753 |
| D_Ko_C04__pb_375_004 | 1388443 | 12,95 | 69,752 | 9,328 | 8,528 |
| S_Up_H47__pb_382_004 | 1561579 | 14,44 | 67,569 | 9,246 | 8,45 |
| S_Up_H29__pb_410_002 | 1472021 | 12,19 | 47,971 | 8,284 | 7,298 |
| E_Vi_C103__pb_410_003 | 1739876 | 12,84 | 49,815 | 7,38 | 6,895 |
| E_Vi_C57__pb_410_004 | 1668218 | 12,98 | 51,044 | 7,778 | 6,812 |
| Pl_Wa_H24__pb_410_006 | 1407040 | 13,19 | 51,352 | 9,371 | 8,795 |
| Pl_Wa_H22__pb_410_005 | 1400972 | 12,92 | 50,659 | 9,222 | 8,814 |
| D_Ko_C13__pb_410_007 | 1449483 | 13,49 | 50,611 | 9,307 | 9,027 |
| S_To_J15__pb_410_010 | 1665168 | 12,16 | 50,03 | 7,301 | 6,787 |
| USA_Wa_B02__pb_410_009 | 1449719 | 12,55 | 47,961 | 8,657 | 8,225 |
| USA_Wa_B01__pb_410_008 | 1461798 | 12,60 | 50,766 | 8,618 | 8,3 |
| S_Up_H24__pb_410_001 | 2155243 | 13,52 | 46,222 | 6,272 | 5,18 |
| S_Up_H32__pb_210_001 | 1599921 | 12,57 | 52,426 | 7,858 | 6,945 |
| S_Up_H32__pb_260_002 | 858882 | 5,71 | 45,288 | 6,644 | 5,916 |
| S_Up_H32__pb_260_003 | 2307285 | 15,06 | 47,199 | 6,527 | 5,744 |
| S_Up_H32__pb_260_001 | 4655884 | 29,74 | 50,116 | 6,388 | 5,647 |
| S_Up_J01__pb_298_001 | 9614265 | 83,33 | 53,658 | 8,667 | 7,923 |
| S_Up_H03__ps_024_006 | 2526698 | 17,22 | 87,77 | 6,814 | 5,6 |
| D_Ko_C36__ps_024_005 | 3684211 | 23,70 | 88,122 | 6,433 | 5,583 |
| D_Ra_C16__ps_024_002 | 2779764 | 19,33 | 98,694 | 6,955 | 6,209 |

| | | | | | |
|---|---|---|---|---|---|
| D_Ko_C15__ps_024_008 | 2851263 | 16,85 | 144,589 | 5,91 | 4,965 |
| S_To_J10__ps_024_010 | 4590885 | 28,05 | 99,725 | 6,109 | 4,886 |
| D_Ko_C31__ps_024_009 | 3888521 | 27,36 | 113,93 | 7,036 | 5,456 |
| S_Up_H37__ps_024_007 | 3248427 | 21,08 | 94,114 | 6,49 | 6,004 |
| S_To_J13__ps_024_019 | 4971825 | 32,39 | 104,698 | 6,513 | 5,659 |
| S_Up_H59__ps_024_022 | 4447600 | 31,70 | 106,717 | 7,127 | 5,921 |
| D_Ra_C05__ps_024_021 | 4463592 | 30,86 | 116,395 | 6,913 | 5,655 |
| E_Vi_C98__ps_038_001 | 2403048 | 16,58 | 77,51 | 6,898 | 5,415 |
| E_Vi_C101__ps_038_004 | 3123591 | 19,89 | 97,839 | 6,368 | 5,098 |
| E_Vi_C100__ps_038_002 | 4011123 | 22,76 | 80,83 | 5,675 | 5,339 |
| RUS_Mp_D06__ps_038_007 | 2106565 | 21,14 | 92,069 | 10,034 | 7,805 |
| RUS_Mp_D08__ps_038_006 | 4114486 | 21,48 | 85,377 | 5,219 | 5,676 |
| RUS_Mp_D04__ps_038_005 | 3524965 | 18,73 | 73,561 | 5,314 | 5,104 |
| Total | 104188441 | 754,80 | 72,57087179 | 7,5347 | 6,719 |

**Table S3.**

Frequencies of individual repeat motifs in filtered insertions and deletions.

| Repeat ID | Frequency | Repeat class | Length |
|---|---|---|---|
| TguERVK7-La_corCor | 1310 | LTR | 670 |
| TguLTRL2-Lc_corCor | 1039 | LTR | 1315 |
| TguERV1-Ld_I_corCor | 838 | LTR | 6022 |
| TguERVL2-La_corCor | 722 | LTR | 564 |
| TguERVL2-Le_corCor | 712 | LTR | 906 |
| A-rich | 646 | Low_complexity | NA |
| (T)n | 620 | Simple_repeat | NA |
| GA-rich | 492 | Low_complexity | NA |
| corCorLTRK23a | 445 | LTR | 294 |
| G-rich | 435 | Low_complexity | NA |
| corCorLTRK15a | 414 | LTR | 966 |
| (A)n | 400 | Simple_repeat | 180 |
| corCorLTRK12a | 379 | LTR | 481 |
| corCorLTRK13a | 349 | LTR | 575 |
| lycPyrLTRL11 | 323 | LTR | 1238 |
| TguLTRL1-La_corCor | 323 | LTR | 640 |
| (TA)n | 258 | Simple_repeat | 180 |
| (C)n | 214 | Simple_repeat | NA |
| corCorLTRK17a_LTR | 211 | LTR | 375 |
| (AT)n | 195 | Simple_repeat | NA |
| corCorLTR1a | 180 | LTR | 461 |
| (G)n | 171 | Simple_repeat | 180 |
| TguLTRL2-La_corCor | 171 | LTR | 1303 |
| corCorLTRK1_I | 163 | LTR | 1736 |
| TguERV1-Lc_LTR_corCor | 159 | LTR | 299 |
| (CCTT)n | 118 | Simple_repeat | NA |
| (GGAA)n | 114 | Simple_repeat | 180 |
| CR1-E1_fAlb | 107 | LINE/CR1 | 437 |
| CR1-J2_Pass | 107 | LINE/CR1 | 4277 |
| (TTCC)n | 104 | Simple_repeat | NA |
| TguLTRL4a | 103 | LTR | 1154 |

**Table S4.**
FST outliers in the all-black German carrion crow and black-and-gray hooded crow comparison based on LR variants.

| Chromosome | Position | Type | Length | FST | Gene downstream | Distance | Gene upstream | Distance |
|---|---|---|---|---|---|---|---|---|
| Super-Scaffold_9_Super-Scaffold_99__chr18 | 10083548 | DEL | 86 | 0.812405 | SLC16A6 | -26679 | ARSG | 301 |
| Sc8eucV_19_HRSCAF_154__chr3 | 78483754 | DEL | 1563 | 0.666196 | GABRR2 | -9425 | UBE2J1 | 850 |
| Sc8eucV_16_HRSCAF_138__chr1 | 112179329 | DEL | 2255 | 0.616612 | NDP | -26270 | EFHC2 | 37138 |
| Sc8eucV_12_HRSCAF_113__chr4A | 10168413 | DEL | 659 | 0.556894 | GRIA3 | -204950 | MCTS1 | 304532 |
| Sc8eucV_10_HRSCAF_100__chr15 | 9876103 | INS | 920 | 0.542418 | SLC5A1 | -25446 | YWHAH | 2969 |
| Sc8eucV_6_HRSCAF_52__chr8 | 81990 | DEL | 1338 | 0.542418 | SLC44A5 | 0 | SLC44A5 | 0 |
| Super-Scaffold_9_Super-Scaffold_99__chr18 | 9080135 | INS | 56 | 0.542418 | LOC104696015 | -966 | BPTF | 15959 |
| Sc8eucV_21_HRSCAF_161__chr4 | 30401968 | DEL | 819 | 0.528082 | GIMD1 | -91886 | DKK2 | 43851 |
| Super-Scaffold_9_Super-Scaffold_99__chr18 | 10160422 | DEL | 608 | 0.515687 | FAM20A | -9970 | LOC104693358 | 21836 |
| Sc8eucV_28_HRSCAF_204__chrZ | 42855849 | DEL | 6523 | 0.498962 | IFT74 | -436763 | CAAP1 | 402328 |
| Sc8eucV_17_HRSCAF_144__chr12 | 14067812 | INS | 134 | 0.498818 | PSMD6 | -25685 | PRICKLE2 | 3640 |
| Sc8eucV_41_HRSCAF_239__chr2 | 11673240 | DEL | 115 | 0.45518 | PITRM1 | -102277 | PFKP | 132560 |
| Sc8eucV_21_HRSCAF_161__chr4 | 10686903 | DEL | 1481 | 0.453426 | MAML3 | -18719 | MGST2 | 32272 |
| Sc8eucV_19_HRSCAF_154__chr3 | 93939333 | INS | 58 | 0.439911 | NA | NA | NA | NA |
| Sc8eucV_28_HRSCAF_204__chrZ | 34154260 | DEL | 567 | 0.437665 | NA | NA | NA | NA |
| Sc8eucV_16_HRSCAF_138__chr1 | 435789 | DEL | 100 | 0.430649 | SLAMF9 | -79446 | LOC104698121 | 31026 |
| Sc8eucV_20_HRSCAF_160__chr1A | 22139103 | DEL | 74 | 0.430649 | PTPRZ1 | 0 | PTPRZ1 | 0 |
| Super-Scaffold_9_Super-Scaffold_99__chr18 | 6551090 | INS | 92 | 0.430649 | CA10 | -972 | UTP18 | 282159 |
| Super-Scaffold_9_Super-Scaffold_99__chr18 | 9954827 | INS | 64 | 0.427794 | AXIN2 | -95524 | RGS9 | 14188 |
| Sc8eucV_10_HRSCAF_100__chr15 | 1130347 | INS | 119 | 0.425766 | GLT1D1 | -31874 | SLC15A4 | 8750 |
| Sc8eucV_14_HRSCAF_135__chr10 | 18758127 | DEL | 70 | 0.425766 | MPI | -16805 | SCAMP2 | 4290 |

| | | | | | | | | |
|---|---|---|---|---|---|---|---|---|
| Sc8eucV_20_HRSCAF_160__chr1A | 40984218 | INS | 295 | 0.425766 | SLC6A15 | -13521 | TSPAN19 | 43707 |
| Sc8eucV_19_HRSCAF_154__chr3 | 64545414 | DEL | 3696 | 0.42464 | MAN1A1 | -326161 | TBC1D32 | 296556 |
| Sc8eucV_19_HRSCAF_154__chr3 | 43076707 | DEL | 569 | 0.417853 | DISC1 | 0 | DISC1 | 0 |
| Sc8eucV_22_HRSCAF_163__chr9 | 24812560 | DEL | 289 | 0.414455 | PCCB | -20580 | PPP2R3A | 13552 |
| Sc8eucV_12_HRSCAF_113__chr4A | 1505992 | DEL | 114 | 0.408223 | LOC104691275 | -372 | EDA2R | 21394 |
| Sc8eucV_5_HRSCAF_38__chr7 | 1789800 | DEL | 221 | 0.408223 | MMADHC | -26020 | LYPD6 | 40702 |
| Sc8eucV_12_HRSCAF_113__chr4A | 16945599 | DEL | 129 | 0.407186 | LOC104691732 | -34917 | PIH1D3 | 21123 |
| Sc8eucV_12_HRSCAF_113__chr4A | 8868927 | DEL | 215 | 0.407186 | NRK | 0 | NRK | 0 |
| Sc8eucV_41_HRSCAF_239__chr2 | 31827203 | DEL | 162 | 0.388603 | ELMO1 | 0 | ELMO1 | 0 |
| Sc8eucV_19_HRSCAF_154__chr3 | 56433950 | DEL | 597 | 0.388519 | ESR1 | 0 | ESR1 | 0 |
| Sc8eucV_2_HRSCAF_5__chr5 | 37341942 | DEL | 400 | 0.387775 | LOC104698601 | 0 | LOC104698601 | 0 |
| Sc8eucV_2_HRSCAF_5__chr5 | 47984643 | DEL | 143 | 0.387279 | TH | -15362 | LOC104687704 | 119783 |
| Sc8eucV_11_HRSCAF_110__chr11 | 2373236 | INS | 131 | 0.381214 | GSE1 | -22151 | GINS2 | 40332 |
| Sc8eucV_6_HRSCAF_52__chr8 | 478781 | DEL | 93 | 0.381214 | NEGR1 | -61593 | ERICH3 | 227018 |
| Super-Scaffold_8_Super-Scaffold_98__chr28 | 1864815 | INS | 149 | 0.381214 | ADAMTS10 | -72880 | ZAP70 | 35464 |
| Sc8eucV_21_HRSCAF_161__chr4 | 41662680 | DEL | 7777 | 0.379839 | SORBS2 | -384955 | MTNR1A | 95511 |
| Sc8eucV_41_HRSCAF_239__chr2 | 47695276 | INS | 146 | 0.379596 | IL6 | -10879 | LOC104689112 | 21839 |
| Sc8eucV_16_HRSCAF_138__chr1 | 2398759 | INS | 68 | 0.37461 | LOC104689368 | -75896 | LOC104689433 | 39839 |
| Sc8eucV_2_HRSCAF_5__chr5 | 38174309 | DEL | 135 | 0.372993 | RTF1 | 0 | RTF1 | 0 |
| Sc8eucV_41_HRSCAF_239__chr2 | 84746331 | INS | 80 | 0.372993 | COBL | -357811 | LOC104685918 | 289065 |
| Sc8eucV_6_HRSCAF_52__chr8 | 1129691 | DEL | 79 | 0.372993 | LOC104695493 | -2511 | WLS | 2023 |
| Sc8eucV_2_HRSCAF_5__chr5 | 15012660 | INS | 75 | 0.371391 | PPP4R4 | 0 | PPP4R4 | 0 |
| Sc8eucV_11_HRSCAF_110__chr11 | 14615394 | INS | 59 | 0.366293 | LOC104687436 | -3416 | POP4 | 358 |
| Sc8eucV_19_HRSCAF_154__chr3 | 54586748 | DEL | 61 | 0.365118 | NOX3 | -113289 | CLDN20 | 189881 |
| Super-Scaffold_8_Super-Scaffold_98__chr28 | 3607722 | DEL | 90 | 0.365118 | LOC104697075 | 0 | LOC104697075 | 0 |
| Super-Scaffold_9_Super-Scaffold_99__chr18 | 10166476 | INS | 151 | 0.365118 | FAM20A | -16024 | LOC104696001 | 23298 |
| Super-Scaffold_9_Super-Scaffold_99__chr18 | 10244723 | DEL | 91 | 0.365118 | LOC104696000 | -7678 | ABCA5 | 1943 |
| Super-Scaffold_143__chr26 | 6291854 | INS | 282 | 0.357143 | PPFIA4 | 0 | PPFIA4 | 0 |

| | | | | | | | | |
|---|---|---|---|---|---|---|---|---|
| Sc8eucV_16_HRSCAF_138__chr1 | 8432823 | INS | 916 | 0.351229 | ROBO1 | -235266 | LOC104689417 | 266974 |
| Sc8eucV_20_HRSCAF_160__chr1A | 27000693 | DEL | 2184 | 0.351229 | IMMP2L | 0 | IMMP2L | 0 |
| Sc8eucV_21_HRSCAF_161__chr4 | 11319706 | INS | 52 | 0.351229 | RNF150 | -822 | LOC104697268 | 95611 |
| Sc8eucV_2_HRSCAF_5__chr5 | 564728 | INS | 100 | 0.351229 | FAM179B | -211063 | KLHL28 | 169095 |
| Sc8eucV_41_HRSCAF_239__chr2 | 9079818 | INS | 460 | 0.351229 | LOC104686440 | 0 | LOC104686440 | 0 |
| Sc8eucV_7_HRSCAF_58__chr13 | 1843425 | DEL | 448 | 0.351229 | LOC104689637 | -6048 | SLC26A2 | 8279 |
| Sc8eucV_16_HRSCAF_138__chr1 | 60691922 | DEL | 912 | 0.34902 | LOC104693136 | -557782 | DIAPH3 | 298389 |
| Sc8eucV_22_HRSCAF_163__chr9 | 24517238 | DEL | 2864 | 0.34902 | MAP3K13 | 0 | MAP3K13 | 0 |
| Super-Scaffold_9_Super-Scaffold_99__chr18 | 5715351 | DEL | 51 | 0.34902 | ACSF2 | 0 | ACSF2 | 0 |
| Sc8eucV_11_HRSCAF_110__chr11 | 2409601 | DEL | 149 | 0.345105 | GSE1 | 0 | GSE1 | 0 |
| Sc8eucV_16_HRSCAF_138__chr1 | 47207238 | INS | 74 | 0.340705 | WASF3 | -4322 | CDK8 | 45576 |
| Sc8eucV_6_HRSCAF_52__chr8 | 23785773 | DEL | 300 | 0.340705 | LOC104695422 | -48461 | CDC73 | 9165 |
| Sc8eucV_41_HRSCAF_239__chr2 | 74069849 | INS | 384 | 0.340533 | CDH9 | 0 | CDH9 | 0 |
| Sc8eucV_12_HRSCAF_113__chr4A | 13659348 | DEL | 54 | 0.339498 | LOC104691679 | 0 | LOC104691679 | 0 |
| Sc8eucV_5_HRSCAF_38__chr7 | 982205 | INS | 92 | 0.337316 | DRC1 | -14449 | GALNT13 | 55917 |
| Sc8eucV_41_HRSCAF_239__chr2 | 101128609 | INS | 324 | 0.336903 | NA | NA | NA | NA |
| Sc8eucV_28_HRSCAF_204__chrZ | 73797802 | INS | 73 | 0.330544 | EDIL3 | -406650 | LOC104695841 | 282404 |
| Sc8eucV_20_HRSCAF_160__chr1A | 67154259 | INS | 51 | 0.329462 | PRR5 | 0 | PRR5 | 0 |
| Sc8eucV_28_HRSCAF_204__chrZ | 13197166 | INS | 573 | 0.329462 | RAI14 | 0 | RAI14 | 0 |
| Sc8eucV_2_HRSCAF_5__chr5 | 5602372 | DEL | 830 | 0.329462 | PRKCH | -64676 | SLC38A6 | 52965 |
| Sc8eucV_6_HRSCAF_52__chr8 | 10448114 | DEL | 60 | 0.329462 | IPO13 | -9585 | ST3GAL3 | 4992 |
| Super-Scaffold_9_Super-Scaffold_99__chr18 | 7026905 | INS | 97 | 0.329462 | LOC104694274 | -17172 | LOC104694356 | 204630 |
| Sc8eucV_2_HRSCAF_5__chr5 | 28446815 | DEL | 53 | 0.326572 | LOC104684460 | -210386 | LOC104684431 | 41038 |
| Super-Scaffold_9_Super-Scaffold_99__chr18 | 2181334 | DEL | 184 | 0.326572 | LOC104695636 | -20399 | LOC104695682 | 1877 |
| Sc8eucV_20_HRSCAF_160__chr1A | 29292175 | INS | 55 | 0.321524 | ADAMTS20 | 0 | ADAMTS20 | 0 |
| Sc8eucV_19_HRSCAF_154__chr3 | 77188793 | DEL | 6950 | 0.321188 | MAP3K7 | -695846 | EPHA7 | 397902 |
| Sc8eucV_2_HRSCAF_5__chr5 | 6552052 | INS | 108 | 0.321188 | WDR89 | 0 | WDR89 | 0 |
| Sc8eucV_41_HRSCAF_239__chr2 | 26678831 | DEL | 1164 | 0.321188 | MRPL32 | -103684 | LOC104688910 | 82671 |

| | | | | | | | | |
|---|---|---|---|---|---|---|---|---|
| Sc8eucV_19_HRSCAF_154__chr3 | 114426297 | DEL | 89 | 0.319956 | DTNB | 0 | DTNB | 0 |
| Sc8eucV_10_HRSCAF_100__chr15 | 11369903 | DEL | 584 | 0.315981 | LOC104694978 | -111902 | AIFM3 | 200211 |
| Super-Scaffold_503__chr27 | 4947702 | DEL | 192 | 0.313759 | LOC104685769 | 0 | LOC104685769 | 0 |
| Sc8eucV_19_HRSCAF_154__chr3 | 66564293 | INS | 160 | 0.312588 | NT5DC1 | -214535 | FRK | 151867 |
| Sc8eucV_1_HRSCAF_4__chr6 | 16956472 | INS | 79 | 0.312588 | KIF20B | 0 | KIF20B | 0 |
| Sc8eucV_21_HRSCAF_161__chr4 | 55571864 | INS | 155 | 0.312588 | PPARGC1A | -48196 | LOC104690704 | 53198 |
| Sc8eucV_20_HRSCAF_160__chr1A | 10564911 | DEL | 621 | 0.311616 | GNAI1 | -144700 | GNAT3 | 241037 |
| Sc8eucV_6_HRSCAF_52__chr8 | 29238870 | DEL | 1209 | 0.311616 | CCDC180 | -528731 | LOC104693596 | 37188 |
| Sc8eucV_5_HRSCAF_38__chr7 | 33426000 | DEL | 7076 | 0.307692 | ZNF804A | -178769 | LOC104698251 | 112821 |
| Sc8eucV_13_HRSCAF_122__chr20 | 1264893 | DEL | 190 | 0.306888 | LOC104698034 | -2824 | LOC104698050 | 42840 |
| Sc8eucV_16_HRSCAF_138__chr1 | 24035795 | DEL | 383 | 0.306888 | LSAMP | -150407 | GAP43 | 481899 |
| Sc8eucV_41_HRSCAF_239__chr2 | 61742801 | DEL | 699 | 0.306143 | CDKAL1 | 0 | CDKAL1 | 0 |
| Sc8eucV_18_HRSCAF_152__chr22 | 4681012 | DEL | 145 | 0.305817 | LOC104693230 | 0 | LOC104693230 | 0 |
| Sc8eucV_19_HRSCAF_154__chr3 | 91662759 | DEL | 122 | 0.305817 | LOC104693575 | -4823 | LOC104693569 | 6609 |
| Sc8eucV_19_HRSCAF_154__chr3 | 59802741 | INS | 52 | 0.3027 | EPB41L2 | -985 | SMLR1 | 118199 |
| Sc8eucV_20_HRSCAF_160__chr1A | 72391930 | DEL | 67 | 0.3027 | CACNA1C | 0 | CACNA1C | 0 |
| Sc8eucV_28_HRSCAF_204__chrZ | 41910148 | DEL | 58 | 0.3027 | SLC12A2 | -121109 | LOC104691976 | 24998 |
| Sc8eucV_28_HRSCAF_204__chrZ | 658358 | INS | 220 | 0.3027 | TCF4 | 0 | TCF4 | 0 |
| Sc8eucV_16_HRSCAF_138__chr1 | 57452531 | DEL | 1336 | 0.301669 | DGKH | -1802 | RGCC | 329040 |
| Sc8eucV_2_HRSCAF_5__chr5 | 45053086 | DEL | 136 | 0.301417 | LRP5 | 0 | LRP5 | 0 |

**Table S5.**

Detailed sample information on sequenced and mapped individuals.

| Genus | Species | Individual ID | Tissue | Sampling location | Sequencing Instrument |
|---|---|---|---|---|---|
| Corvus | brachyrhynchos | USA_CA_B01 | tissue, blood | USA, California, Shasty County | Illumina HiSeq2000 |
| Corvus | brachyrhynchos | USA_CA_B03 | tissue, blood | USA, California, Shasta, Cottonwood | Illumina HiSeq2001 |
| Corvus | brachyrhynchos | USA_CA_B08 | unknown | USA, California | Illumina HiSeq2002 |
| Corvus | brachyrhynchos | USA_NJ_B02 | tissue, blood | USA, New Jersey, Union County, Westfield | Illumina HiSeq2003 |
| Corvus | brachyrhynchos | USA_NY_B03 | tissue, blood | USA, New York, Suffolk County, Northport | Illumina HiSeq2004 |
| Corvus | brachyrhynchos | USA_NY_B04 | tissue, blood | USA, New York, Nassau County, Wantagh | Illumina HiSeq2005 |
| Corvus | corone cornix | B_So_H01 | blood | Bulgaria, Sofia | Illumina HiSeq2006 |
| Corvus | corone cornix | B_So_H02 | blood | Bulgaria, Sofia | Illumina HiSeq2007 |
| Corvus | corone cornix | B_So_H03 | blood | Bulgaria, Sofia | Illumina HiSeq2008 |
| Corvus | corone cornix | B_So_H04 | blood | Bulgaria, Sofia | Illumina HiSeq2009 |
| Corvus | corone cornix | B_SZ_H01 | blood | Bulgaria, Stora Zagora | Illumina HiSeq2010 |
| Corvus | corone cornix | B_SZ_H02 | blood | Bulgaria, Stora Zagora | Illumina HiSeq2011 |
| Corvus | corone cornix | B_un_H01 | blood | Bulgaria | Illumina HiSeq2012 |
| Corvus | corone cornix | ISR_TA_H01 | blood | Israel, Tel Aviv | Illumina HiSeq2013 |
| Corvus | corone cornix | ISR_TA_H02 | blood | Israel, Tel Aviv | Illumina HiSeq2014 |
| Corvus | corone cornix | ISR_TA_H04 | blood | Israel, Tel Aviv | Illumina HiSeq2015 |
| Corvus | corone cornix | ITA_Ro_H01 | blood | Italy, Rome | Illumina HiSeq2016 |
| Corvus | corone cornix | ITA_Ro_H02 | blood | Italy, Rome | Illumina HiSeq2017 |
| Corvus | corone cornix | ITA_Ro_H03 | blood | Italy, Rome | Illumina HiSeq2018 |
| Corvus | corone cornix | ITA_Ro_H04 | blood | Italy, Rome | Illumina HiSeq2019 |
| Corvus | corone cornix | ITA_Ro_H05 | blood | Italy, Rome | Illumina HiSeq2020 |
| Corvus | corone cornix | ITA_Ro_H06 | blood | Italy, Rome | Illumina HiSeq2021 |
| Corvus | corone cornix | ITA_Ro_H07 | blood | Italy, Rome | Illumina HiSeq2022 |
| Corvus | corone cornix | ITA_Ro_H08 | blood | Italy, Rome | Illumina HiSeq2023 |
| Corvus | corone cornix | ITA_Ro_H09 | tissue | Italy, Rome | Illumina HiSeq2024 |

| | | | | | |
|---|---|---|---|---|---|
| Corvus | corone cornix | ITA_Ro_H10 | tissue | Italy, Rome | Illumina HiSeq2025 |
| Corvus | corone cornix | ITA_Ro_H11 | tissue | Italy, Rome | Illumina HiSeq2026 |
| Corvus | corone cornix | ITA_Ro_H12 | tissue | Italy, Rome | Illumina HiSeq2027 |
| Corvus | corone cornix | ITA_Ro_H13 | tissue | Italy, Rome | Illumina HiSeq2028 |
| Corvus | corone cornix | ITA_Ro_H14 | tissue | Italy, Rome | Illumina HiSeq2029 |
| Corvus | corone cornix | PL_Wa_H02 | blood | Poland, Warsaw | Illumina HiSeq2030 |
| Corvus | corone cornix | PL_Wa_H03 | blood | Poland, Warsaw | Illumina HiSeq2031 |
| Corvus | corone cornix | PL_Wa_H05 | blood | Poland, Warsaw | Illumina HiSeq2032 |
| Corvus | corone cornix | PL_Wa_H06 | blood | Poland, Warsaw | Illumina HiSeq2033 |
| Corvus | corone cornix | PL_Wa_H09 | blood | Poland, Warsaw | Illumina HiSeq2034 |
| Corvus | corone cornix | PL_Wa_H11 | blood | Poland, Warsaw | Illumina HiSeq2035 |
| Corvus | corone cornix | PL_Wa_H14 | blood | Poland, Warsaw | Illumina HiSeq2036 |
| Corvus | corone cornix | PL_Wa_H16 | blood | Poland, Warsaw | Illumina HiSeq2037 |
| Corvus | corone cornix | PL_Wa_H17 | blood | Poland, Warsaw | Illumina HiSeq2038 |
| Corvus | corone cornix | PL_Wa_H22 | blood | Poland, Warsaw | Illumina HiSeq2039 |
| Corvus | corone cornix | PL_Wa_H23 | blood | Poland, Warsaw | Illumina HiSeq2040 |
| Corvus | corone cornix | PL_Wa_H32 | blood | Poland, Warsaw | Illumina HiSeq2041 |
| Corvus | corone cornix | PL_Wa_H35 | blood | Poland, Warsaw | Illumina HiSeq2042 |
| Corvus | corone cornix | PL_Wa_H50 | blood | Poland, Warsaw | Illumina HiSeq2043 |
| Corvus | corone cornix | PL_Wa_H52 | blood | Poland, Warsaw | Illumina HiSeq2044 |
| Corvus | corone cornix | RUS_Ki_H02 | blood | Russia, Kirov | Illumina HiSeq2045 |
| Corvus | corone cornix | RUS_Ki_H03 | blood | Russia, Kirov | Illumina HiSeq2046 |
| Corvus | corone cornix | RUS_Ki_H04 | blood | Russia, Kirov | Illumina HiSeq2047 |
| Corvus | corone cornix | RUS_No_H02 | blood | Russia, Novosibirsk | Illumina HiSeq2048 |
| Corvus | corone cornix | RUS_No_H03 | blood | Russia, Novosibirsk | Illumina HiSeq2049 |
| Corvus | corone cornix | RUS_Tu_H01 | blood | Russia, Tyumen | Illumina HiSeq2050 |
| Corvus | corone cornix | S_Ri_H05 | liver | Sweden, Rimbo | Illumina HiSeq2051 |
| Corvus | corone cornix | S_Ri_H07 | liver | Sweden, Rimbo | Illumina HiSeq2052 |
| Corvus | corone cornix | S_Ri_H23 | liver | Sweden, Rimbo | Illumina HiSeq2053 |

| | | | | | |
|---|---|---|---|---|---|
| Corvus | corone cornix | S_Ri_H29 | liver | Sweden, Rimbo | Illumina HiSeq2054 |
| Corvus | corone cornix | S_Ri_H43 | liver | Sweden, Rimbo | Illumina HiSeq2055 |
| Corvus | corone cornix | S_Up_H03 | blood | Sweden, Uppsala | Illumina HiSeq2056 |
| Corvus | corone cornix | S_Up_H09 | blood | Sweden, Uppsala | Illumina HiSeq2057 |
| Corvus | corone cornix | S_Up_H16 | blood | Sweden, Uppsala | Illumina HiSeq2058 |
| Corvus | corone cornix | S_Up_H24 | blood | Sweden, Uppsala | Illumina HiSeq2059 |
| Corvus | corone cornix | S_Up_H29 | blood | Sweden, Uppsala | Illumina HiSeq2060 |
| Corvus | corone cornix | S_Up_H37 | blood | Sweden, Uppsala | Illumina HiSeq2061 |
| Corvus | corone cornix | S_Up_H43 | blood | Sweden, Uppsala | Illumina HiSeq2062 |
| Corvus | corone cornix | S_Up_H47 | blood | Sweden, Uppsala | Illumina HiSeq2063 |
| Corvus | corone cornix | S_Up_H51 | blood | Sweden, Uppsala | Illumina HiSeq2064 |
| Corvus | corone cornix | S_Up_H52 | blood | Sweden, Uppsala | Illumina HiSeq2065 |
| Corvus | corone corone | D_Ko_C02 | blood | Germany, Konstanz | Illumina HiSeq2066 |
| Corvus | corone corone | D_Ko_C04 | blood | Germany, Konstanz | Illumina HiSeq2067 |
| Corvus | corone corone | D_Ko_C05 | blood | Germany, Konstanz | Illumina HiSeq2068 |
| Corvus | corone corone | D_Ko_C08 | blood | Germany, Konstanz | Illumina HiSeq2069 |
| Corvus | corone corone | D_Ko_C11 | blood | Germany, Konstanz | Illumina HiSeq2070 |
| Corvus | corone corone | D_Ko_C13 | blood | Germany, Konstanz | Illumina HiSeq2071 |
| Corvus | corone corone | D_Ko_C15 | blood | Germany, Konstanz | Illumina HiSeq2072 |
| Corvus | corone corone | D_Ko_C19 | blood | Germany, Konstanz | Illumina HiSeq2073 |
| Corvus | corone corone | D_Ko_C20 | blood | Germany, Konstanz | Illumina HiSeq2074 |
| Corvus | corone corone | D_Ra_C01 | blood | Germany, Radolfzell | Illumina HiSeq2075 |
| Corvus | corone corone | D_Ra_C05 | blood | Germany, Radolfzell | Illumina HiSeq2076 |
| Corvus | corone corone | D_Ra_C06 | blood | Germany, Radolfzell | Illumina HiSeq2077 |
| Corvus | corone corone | D_Ra_C11 | blood | Germany, Radolfzell | Illumina HiSeq2078 |
| Corvus | corone corone | D_Ra_C14 | blood | Germany, Radolfzell | Illumina HiSeq2079 |
| Corvus | corone corone | D_Ra_C16 | blood | Germany, Radolfzell | Illumina HiSeq2080 |
| Corvus | corone corone | E_Vi_C01 | blood | Spain, Villaseca de la Sorriba | Illumina HiSeq2081 |
| Corvus | corone corone | E_Vi_C05 | blood | Spain, Villaseca de la Sorriba | Illumina HiSeq2082 |

| | | | | | |
|---|---|---|---|---|---|
| Corvus | corone corone | E_Vi_C08 | blood | Spain, Villaseca de la Sorriba | Illumina HiSeq2083 |
| Corvus | corone corone | E_Vi_C14 | blood | Spain, Villaseca de la Sorriba | Illumina HiSeq2084 |
| Corvus | corone corone | E_Vi_C19 | blood | Spain, Villaseca de la Sorriba | Illumina HiSeq2085 |
| Corvus | corone corone | E_Vi_C22 | blood | Spain, Villaseca de la Sorriba | Illumina HiSeq2086 |
| Corvus | corone corone | E_Vi_C23 | blood | Spain, Villaseca de la Sorriba | Illumina HiSeq2087 |
| Corvus | corone corone | E_Vi_C32 | blood | Spain, Villaseca de la Sorriba | Illumina HiSeq2088 |
| Corvus | corone corone | E_Vi_C37 | blood | Spain, Villaseca de la Sorriba | Illumina HiSeq2089 |
| Corvus | corone corone | E_Vi_C44 | blood | Spain, Villaseca de la Sorriba | Illumina HiSeq2090 |
| Corvus | corone corone | E_Vi_C46 | blood | Spain, Villaseca de la Sorriba | Illumina HiSeq2091 |
| Corvus | corone corone | E_Vi_C48 | blood | Spain, Villaseca de la Sorriba | Illumina HiSeq2092 |
| Corvus | corone corone | E_Vi_C51 | blood | Spain, Villaseca de la Sorriba | Illumina HiSeq2093 |
| Corvus | corone corone | E_Vi_C57 | blood | Spain, Villaseca de la Sorriba | Illumina HiSeq2094 |
| Corvus | corone corone | E_Vi_C58 | blood | Spain, Villaseca de la Sorriba | Illumina HiSeq2095 |
| Corvus | corone coroneXcorone cornix | IRL_Lm_H07 | blood | Ireland, Lough Money Farm | Illumina HiSeq2096 |
| Corvus | corone coroneXcorone cornix | IRL_Lm_H08 | blood | Ireland, Lough Money Farm | Illumina HiSeq2097 |
| Corvus | corone coroneXcorone cornix | IRL_Lm_H10 | blood | Ireland, Lough Money Farm | Illumina HiSeq2098 |
| Corvus | corone coroneXcorone cornix | IRL_Lm_H12 | blood | Ireland, Lough Money Farm | Illumina HiSeq2099 |
| Corvus | corone coroneXcorone cornix | IRL_Lm_H15 | blood | Ireland, Lough Money Farm | Illumina HiSeq2100 |
| Corvus | corone coroneXcorone cornix | IRL_Lm_H16 | blood | Ireland, Lough Money Farm | Illumina HiSeq2101 |
| Corvus | corone coroneXcorone cornix | RUS_Ke_Y01 | blood | Russia, Kemerovo | Illumina HiSeq2102 |
| Corvus | corone coroneXcorone cornix | RUS_Ke_Y02 | blood | Russia, Kemerovo | Illumina HiSeq2103 |
| Corvus | corone coroneXcorone cornix | RUS_Ke_Y03 | blood | Russia, Kemerovo | Illumina HiSeq2104 |
| Corvus | corone coroneXcorone cornix | RUS_Ke_Y05 | blood | Russia, Kemerovo | Illumina HiSeq2105 |
| Corvus | corone coroneXcorone cornix | RUS_Ke_Y06 | blood | Russia, Kemerovo | Illumina HiSeq2106 |
| Corvus | corone orientalis | RUS_Kr_O01 | blood | Russia, Krasnoyarsky | Illumina HiSeq2107 |
| Corvus | corone orientalis | RUS_Kr_O02 | blood | Russia, Krasnoyarsky | Illumina HiSeq2108 |

| | | | | | |
|---|---|---|---|---|---|
| Corvus | corone orientalis | RUS_Kr_O03 | blood | Russia, Krasnoyarsky | Illumina HiSeq2109 |
| Corvus | corone orientalis | RUS_Kr_O04 | blood | Russia, Krasnoyarsky | Illumina HiSeq2110 |
| Corvus | corone orientalis | RUS_Pr_O01 | tissue, blood | Russia, Primorsky | Illumina HiSeq2111 |
| Corvus | corone orientalis | RUS_Pr_O02 | tissue, blood | Russia, Primorsky | Illumina HiSeq2112 |
| Corvus | corone orientalis | RUS_Pr_O03 | tissue, blood | Russia, Primorsky | Illumina HiSeq2113 |
| Corvus | corone orientalis | RUS_Pr_O04 | tissue, blood | Russia, Primorsky | Illumina HiSeq2114 |
| Corvus | corone orientalis | RUS_Pr_O05 | tissue, blood | Russia, Primorsky | Illumina HiSeq2115 |
| Corvus | corone orientalis | RUS_Tv_O01 | blood | Russia, Tuva | Illumina HiSeq2116 |
| Corvus | corone orientalis | RUS_Tv_O02 | blood | Russia, Tuva | Illumina HiSeq2117 |
| Corvus | corone orientalis | RUS_Ya_O01 | tissue | Russia, Yakutsk | Illumina HiSeq2118 |
| Corvus | corone orientalis | RUS_Ya_O02 | blood | Russia, Yakutsk | Illumina HiSeq2119 |
| Corvus | corone orientalis | RUS_Ya_O03 | blood | Russia, Yakutsk | Illumina HiSeq2120 |
| Corvus | dauuricus | CHN_Gu_D02 | liver | China, Guangxi | Illumina HiSeq2121 |
| Corvus | dauuricus | MON_Kh_D01 | blood | Mongolia, Khentii | Illumina HiSeq2122 |
| Corvus | dauuricus | MON_Kh_D02 | blood | Mongolia, Khentii | Illumina HiSeq2123 |
| Corvus | dauuricus | MON_Kh_D03 | blood | Mongolia, Khentii | Illumina HiSeq2124 |
| Corvus | monedula | S_Ri_J01 | liver | Sweden, Rimbo | Illumina HiSeq2125 |
| Corvus | monedula | S_Ri_J02 | liver | Sweden, Rimbo | Illumina HiSeq2126 |
| Corvus | monedula | S_Ri_J03 | liver | Sweden, Rimbo | Illumina HiSeq2127 |
| Corvus | monedula | S_Ri_J08 | liver | Sweden, Rimbo | Illumina HiSeq2128 |
| Corvus | pectoralis/torquatus | CHN_Gu_P01 | tissue | China, Guangxi | Illumina HiSeq2129 |
| Corvus | pectoralis/torquatus | Un_un_P01 | tissue | unknown | Illumina HiSeq2130 |
| Corvus | pectoralis/torquatus | Un_un_P02 | tissue | unknown | Illumina HiSeq2131 |
| Corvus | corone cornix | S_Up_H32 | blood | Sweden,Uppsala | PacBio RSII |
| Corvus | corone cornix | S_Up_H03 | blood | Sweden,Uppsala | PacBio Sequel |
| Corvus | corone cornix | S_Up_H47 | blood | Sweden,Uppsala | PacBio RSII |
| Corvus | corone cornix | S_Up_H37 | blood | Sweden,Uppsala | PacBio Sequel |
| Corvus | corone cornix | S_Up_H24 | blood | Sweden,Uppsala | PacBio RSII |
| Corvus | corone cornix | S_Up_H29 | blood | Sweden,Uppsala | PacBio RSII |

| | | | | | |
|---|---|---|---|---|---|
| Corvus | corone cornix | S_Up_H59 | blood | Sweden,Uppsala | PacBio Sequel |
| Corvus | corone cornix | Pl_Wa_H22 | blood | Poland,Warsaw | PacBio RSII |
| Corvus | corone cornix | Pl_Wa_H24 | blood | Poland,Warsaw | PacBio RSII |
| Corvus | corone corone | D_Ko_C04 | blood | Germany,Konstanz | PacBio RSII |
| Corvus | corone corone | D_Ko_C13 | blood | Germany,Konstanz | PacBio RSII |
| Corvus | corone corone | D_Ko_C15 | blood | Germany,Konstanz | PacBio Sequel |
| Corvus | corone corone | D_Ko_C29 | blood | Germany,Konstanz | PacBio RSII |
| Corvus | corone corone | D_Ra_C16 | blood | Germany,Radolfzell | PacBio Sequel |
| Corvus | corone corone | D_Ko_C36 | blood | Germany,Konstanz | PacBio Sequel |
| Corvus | corone corone | D_Ra_C05 | blood | Germany,Radolfzell | PacBio Sequel |
| Corvus | corone corone | D_Ko_C31 | blood | Germany,Konstanz | PacBio Sequel |
| Corvus | corone corone | E_Vi_C57 | blood | Spain,La Sorriba | PacBio RSII |
| Corvus | corone corone | E_Vi_C58 | blood | Spain,La Sorriba | PacBio RSII |
| Corvus | corone corone | E_Vi_C98 | blood | Spain,La Sorriba | PacBio Sequel |
| Corvus | corone corone | E_Vi_C100 | blood | Spain,La Sorriba | PacBio Sequel |
| Corvus | corone corone | E_Vi_C101 | blood | Spain,La Sorriba | PacBio Sequel |
| Corvus | corone corone | E_Vi_C103 | blood | Spain,La Sorriba | PacBio RSII |
| Corvus | brachyrhynchos | USA_Wa_B01 | blood | USA,Seattle | PacBio RSII |
| Corvus | brachyrhynchos | USA_Wa_B02 | blood | USA,Seattle | PacBio RSII |
| Corvus | monedula | S_Up_J01 | blood | Sweden,Uppsala | PacBio RSII |
| Corvus | monedula | S_To_J13 | blood | Sweden,Aspa | PacBio Sequel |
| Corvus | monedula | S_To_J10 | blood | Sweden,Aspa | PacBio Sequel |
| Corvus | monedula | S_To_J14 | blood | Sweden,Aspa | PacBio RSII |
| Corvus | monedula | S_To_J15 | blood | Sweden,Aspa | PacBio RSII |
| Corvus | dauuricus | RUS_Mp_D04 | blood | Russia,Muraviovka Park | PacBio Sequel |
| Corvus | dauuricus | RUS_Mp_D06 | blood | Russia,Muraviovka Park | PacBio Sequel |
| Corvus | dauuricus | RUS_Mp_D08 | blood | Russia,Muraviovka Park | PacBio Sequel |
| Corvus | corone corone | D_Ra_C05 | blood | Germany, Radolfzell | BioNano Irys |
| Corvus | corone corone | D_Ko_C04 | blood | Germany, Konstanz | BioNano Irys |

| | | | | | |
|---|---|---|---|---|---|
| Corvus | corone corone | D_Ko_C13 | blood | Germany, Konstanz | BioNano Irys |
| Corvus | corone corone | D_Ko_C29 | blood | Germany, Konstanz | BioNano Irys |
| Corvus | corone corone | D_Ko_C36 | blood | Germany, Konstanz | BioNano Irys |
| Corvus | corone corone | E_Vi_C101 | blood | Spain,La Sorriba | BioNano Irys |
| Corvus | corone corone | E_Vi_C103 | blood | Spain,La Sorriba | BioNano Irys |
| Corvus | corone corone | E_Vi_C57 | blood | Spain,La Sorriba | BioNano Irys |
| Corvus | corone corone | E_Vi_C58 | blood | Spain,La Sorriba | BioNano Irys |
| Corvus | monedula | S_To_J17 | blood | Sweden, Tovetorp | BioNano Irys |
| Corvus | corone cornix | S_Up_H03 | blood | Sweden, Uppsala | BioNano Irys |
| Corvus | corone cornix | S_Up_H29 | blood | Sweden, Uppsala | BioNano Irys |
| Corvus | corone cornix | S_Up_H32 | blood | Sweden, Uppsala | BioNano Irys |
| Corvus | corone cornix | S_Up_H37 | blood | Sweden, Uppsala | BioNano Irys |
| Corvus | corone cornix | S_Up_H59 | blood | Sweden, Uppsala | BioNano Irys |
| Corvus | monedula | S_Up_J01 | blood | Sweden, Uppsala | BioNano Irys |

**Table S6.**

Statistics of optical maps and map assemblies.

| Sample | data over 150 Kbp | N50 (Mbp) | Maprate to *Corvus cornix* assembly | N50 | genome size | Eff coverage |
|---|---|---|---|---|---|---|
| D_Ra_C05 | 313.7037 Gbp | 0.2272 | 52.40% | 0.759 | 1202.661 | 106.71 |
| E_Vi_C58 | 149.5706 Gbp | 0.2111 | 44.00% | 0.755 | 1135.633 | 44.05 |
| S_Up_H59 | 132.0199 Gbp | 0.2109 | 50.60% | 0.798 | 1136.396 | 75.20 |
| D_Ko_C29 | 113.2390 Gbp | 0.2164 | 45.70% | 0.647 | 1154.888 | 47.49 |
| E_Vi_C101 | 221.5878 Gbp | 0.2566 | 71.30% | 1.064 | 1306.967 | 130.50 |
| S_To_J17 | 200.8957 Gbp | 0.2336 | 36.40% | 0.882 | 1309.932 | 94.89 |
| S_Up_H32 | 107.2551 Gbp | 0.2171 | 67.20% | 0.763 | 1108.333 | 59.74 |
| S_Up_J01 | 188.0292 Gbp | 0.2438 | 29.30% | 0.812 | 1201.969 | 73.28 |
| D_Ko_C04 | 58 Gbp | 0.1426 | 49.70% | 0.223 | 529.43 | 19.89 |
| D_Ko_C13 | 48.0589 Gbp | 0.1978 | 54.30% | 0.4 | 1291.57 | 32.78 |
| D_Ko_C36 | 141.6023 Gbp | 0.2278 | 69.00% | 1.044 | 1236.42 | 82.23 |
| E_Vi_C103 | 194.30 Gbp | 0.2516 | 65.20% | 0.848 | 1207.24 | 93.00 |
| E_Vi_C57 | 356.49 Gbp | 0.2167 | 55.80% | 0.726 | 1449.28 | 166.50 |
| S_Up_H03 | 178.58 Gbp | 0.2311 | 66.50% | 0.826 | 1192.34 | 89.78 |
| S_Up_H29 | 141.49 Gbp | 0.2488 | 70.70% | 0.824 | 1203.84 | 85.12 |
| S_Up_H37 | 166.40 Gbp | 0.2411 | 68.40% | 0.829 | 1216.61 | 99.61 |
